# Three Dimensional Multiscalar Neurovascular Nephron Connectivity Map of the Human Kidney Across the Lifespan

**DOI:** 10.1101/2024.07.29.605633

**Authors:** Liam McLaughlin, Bo Zhang, Siddharth Sharma, Amanda L. Knoten, Madhurima Kaushal, Jeffrey M. Purkerson, Heidy Huyck, Gloria S. Pryhuber, Joseph P. Gaut, Sanjay Jain

## Abstract

The human kidney is a vital organ with a remarkable ability to coordinate the activity of up to a million nephrons, its main functional tissue unit (FTU), and maintain homeostasis. We developed tissue processing and analytical methods to construct a 3D map of neurovascular nephron connectivity of the human kidney and glean insights into how this structural organization enables coordination of various functions of the nephron, such as glomerular filtration, solute and water absorption, secretion by the tubules, and regulation of blood flow and pressure by the juxtaglomerular apparatus, in addition to how these functions change across disease and lifespans. Using light sheet fluorescence microscopy (LSFM) and morphometric analysis we discovered changes in anatomical orientation of the vascular pole, glomerular density, volume, and innervation through postnatal development and ageing. The extensive nerve network exists from cortex FTUs to medullary loop of Henle, providing connectivity within segments of the same nephron, and between separate nephrons. The nerves organize glomeruli into discreet communities (in the same network of nerves). Adjacent glomerular communities are connected to intercommunal “mother glomeruli” by nerves, a pattern repeating throughout the cortex. These neuro-nephron networks are not developed in postnatal kidneys and are disrupted in diseased kidneys (diabetic or hydronephrosis). This structural organization likely poises the entire glomerular and juxtaglomerular FTUs to synchronize responses to perturbations in fluid homeostasis, utilizing mother glomeruli as network control centers.

## Introduction

The functions of organs and tissues are dependent on precise structural organization that begins during development and is maintained in adulthood. This is critical for vital organs such as the human kidney where there is a complex organization of the main functional tissue unit (FTU), the nephron, across a cortex and medulla in association with surrounding neurovascular networks to maintain fluid homeostasis, blood pressure and other endocrine activities ^1^. Various parts of the nephron and their associations with surrounding structures have distinct functions including filtration by the glomerulus, regulation of blood flow and pressure by the juxtaglomerular apparatus, absorption and secretion of various solutes by the tubules, and water balance by the collecting system facilitated by a complex array of cell types ^2, 3^. These essential roles in fluid homeostasis and blood pressure regulation also render various nephron segments as targets for medications to control fluid balance and blood pressure ^4^. Model system studies have demonstrated complex positive and negative feedback loops between renal sensory and sympathetic nerves through their connections with different parts of the nephron to regulate renal blood flow and salt balance and in kidney dysfunction renal denervation can alleviate fibrosis ^5^. Developmental studies have begun to show evidence that sensory and sympathetic nerves follow the blood vessel in the kidney at a macroscopic level to their respective vascular poles of the glomeruli ^6^. There at the juxtaglomerular apparatus (JGA), macula densa (MD) cells of the distal nephron are sensitive to varying sodium concentrations in the filtrate due to proximal tubular dysfunction and can modulate the activity of the renin angiotensin system (RAAS) through a complex interaction between sensory or sympathetic nerves and Renin producing granular cell to regulate glomerular hemodynamics. However, it is unclear how millions of nephrons coordinate these individual responses from each nephron in the human kidney, nor what anatomical and structural patterning is necessary to enable maturity of kidney function from postnatal to adult kidneys and how this organization changes with declining kidney function with age and disease.

To better understand the structural connectivity of neurovasculature with the nephron segments and how FTUs of the nephron change across lifespan and in disease, we developed 3D multiplex immunofluorescence methods and analytical pipelines using light sheet fluorescence microscopy (LSFM) on thick human kidney slices across the cortical-medullary axis and at various ages for the Human BioMolecular Atlas Program (HuBMAP) ^7^. We report that glomerular attain a highly polarized orientation with increasing nerve density around them as development progresses. They organize into communities that are interconnected through nerves. The 3D image analyses revealed both sensory and sympathetic nerves contribute to the neural networks. The afferent/efferent nerves only distinctly diverge beneath the MD, which showed only sympathetic innervation at the JGA, but had sensory innervation along the MD’s flanks. Neural connectivity was observed within multiple segments of the same nephron, such as from the glomerulus to its thick ascending limb (TAL) and proximal convoluted tubule (PCT), between glomeruli, and between glomeruli and segments of other nephrons. Importantly, the glomerular communities showed several patterns of innervation and most notably, demonstrated adjacent glomerular communities connected via inter-communal glomeruli that we deemed “mother gloms,” based on the concept of highly connected mother tree hub points^8^￼. The communities were interconnected with mother glomeruli throughout the cortex in a repetitive pattern thus providing a structural organization that is poised to sense or relay signals throughout the nephron and that we hypothesize is engineered to coordinate physiological responses by decreasing burden on individual nephrons. The network was severely diminished in diabetic patients or obstructive nephropathy samples with advanced kidney disease suggesting a visceral nephropathy in chronic kidney disease.

## RESULTS

### Overall strategy for 3D LSFM

Architecturally complex solid tissue such as kidney posit challenges for 3D volumetric imaging, impacting tissue penetration of reagents, clearing methods, imaging, and analysis. We tested several tissue clearing methods including passive CLARITY ^9^, iDISCO ^10^, CUBIC ^11^, and CLARITY SHIELD ^12^ on fixed thick tissue slices from nephrectomies or deceased donors with CLARITY SHIELD as the method of choice here (**Figure 1** and data not shown). Our goal was to define anatomical organization of select functional tissue units (FTUs) along the nephron in relation to neuronal patterning, morphology at macro and microscopic scales and across various ages ranging from neonatal to aged individuals and in some cases samples with disease **(Supplemental Table S1).** We examined the overall organization at low power (5X) and then select field of views (FOVs) in 3D at high power (20X) with depth ranging from 1-3mm. In general, kidney structures / cell types evaluated in 3D included nerves, vessels, glomeruli (gloms, podocytes immunostained for NPHS1 or PODXL), collecting duct (CD, principal cells immunostained for AQP2), proximal tubules (PT, PT cells immunostained for LRP2) and thick ascending limb (TAL, TAL cell immunostained for UMOD) of the loop of Henle (LOH)) (**Supplemental Table S2)**. For nerves, TUJ1 (targeting TUBB3) as a pan neuronal marker, and for sensory and sympathetic nerves CGRP and TH, respectively, were used. For endothelial cells in the blood vessels CD31 or CD34 or PODXL were used. In total, 23 samples across 15 patients were imaged using LSFM (**Supplemental Table S1)** and 22 samples analyzed for this study. A summary of the pipeline is displayed in **Figure 1**.

**Figure 1.**
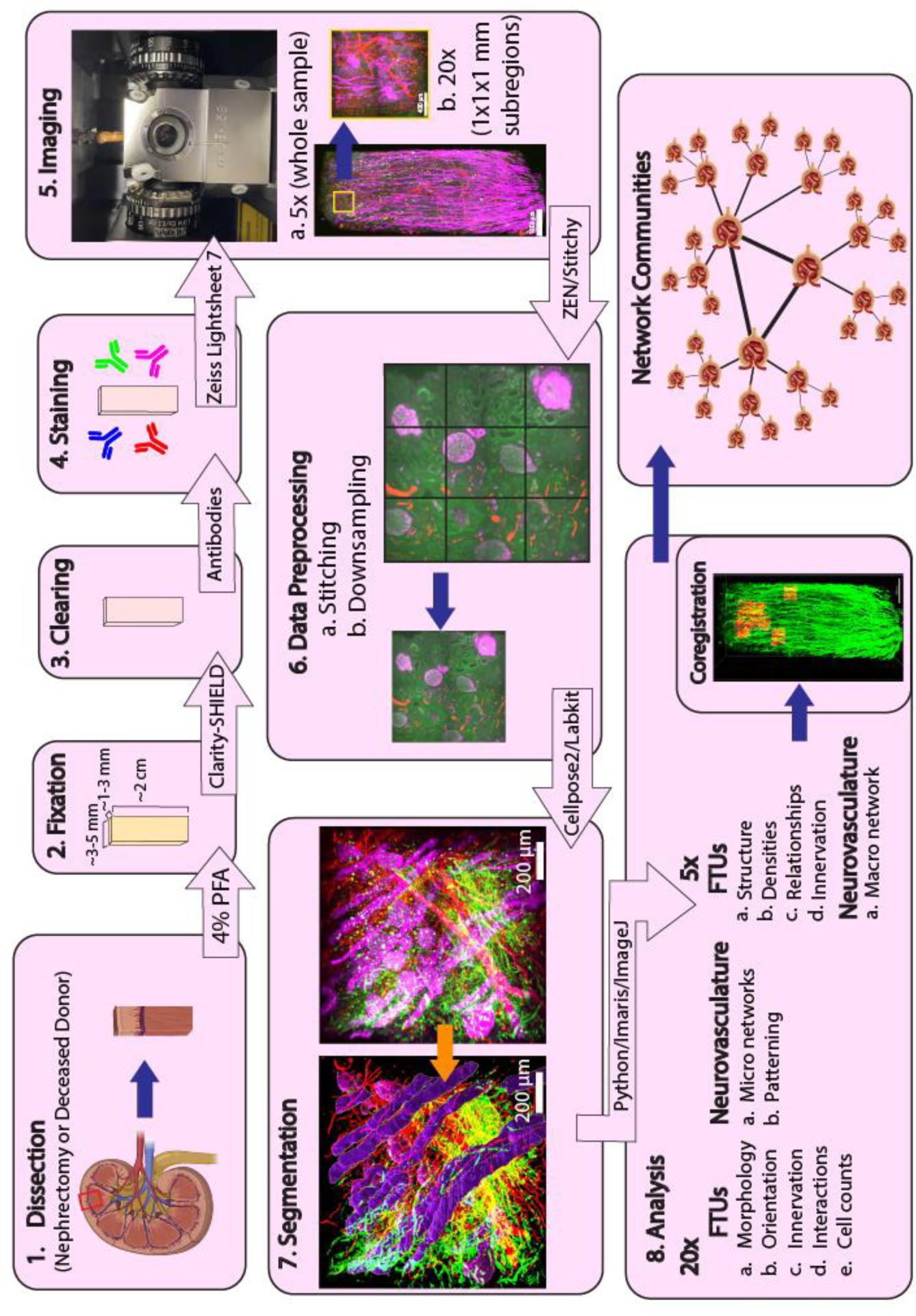
End-to-end pipeline for light sheet fluorescence microscopy from tissue processing to imaging to analysis. 1-4, Tissue processing and immunofluorescence staining of corticomedullary strips of tissue slices from kidney, preserving in fixative, active clearing using and performing multiplex immunofluorescence staining with antibodies targeting key functional tissue units of the kidney and / or neurovasculature. The staining was typically done for 5 targets across the Cy5, Cy3, 488 and 405 channels. 5, LSFM imaging. Cleared and post IF samples were imaged using Zeiss LSFM to obtain gross macroscopic view of the entire specimen at 5X followed by a series of high-resolution, 20x images to visualize subregions along cortico-medullary axis. These 20x images were typically 1x1x1 mm, although a small number were larger. 6, Data processing. Prior to any analysis, data was stitched and downsampled by 4x in both X and Y using either ZEN Blue 3.1 software or Stitchy. 7, Data were segmented for structures of interest utilizing either Cellpose or Labkit. 8, Qualitative and quantitative data analysis for various structures and their relationships was accomplished using commercial software such as Imaris or ImageJ, in addition to in-house Python code. This included coregistration of 20x subregions onto 5x volumes, and algorithmic extraction of neural networks between glomeruli.

### Morphological organization of neurovasculature in relation to FTUs in the cortico-medullary axis

Much of our knowledge about kidney organization comes from 2D sections that are limited in providing relationships between different nephron segments and extracellular material, including their connectivity. To gain insights into structure-function beyond 3D vascular networks, we first evaluated neural network in relation to human kidney FTUs. The 3D volumetric imaging using LSFM with the indicated markers from a region of the kidney lobe along the corticomedullary axis revealed dense innervation of the cortex and medulla, with nerves following the medium and small sized vessels in the cortex and medulla, alongside successful visualization of all labeled structures and their 3D relationships (**Figure 2a, supplementary movie 1, 5x**).

**Figure 2.**
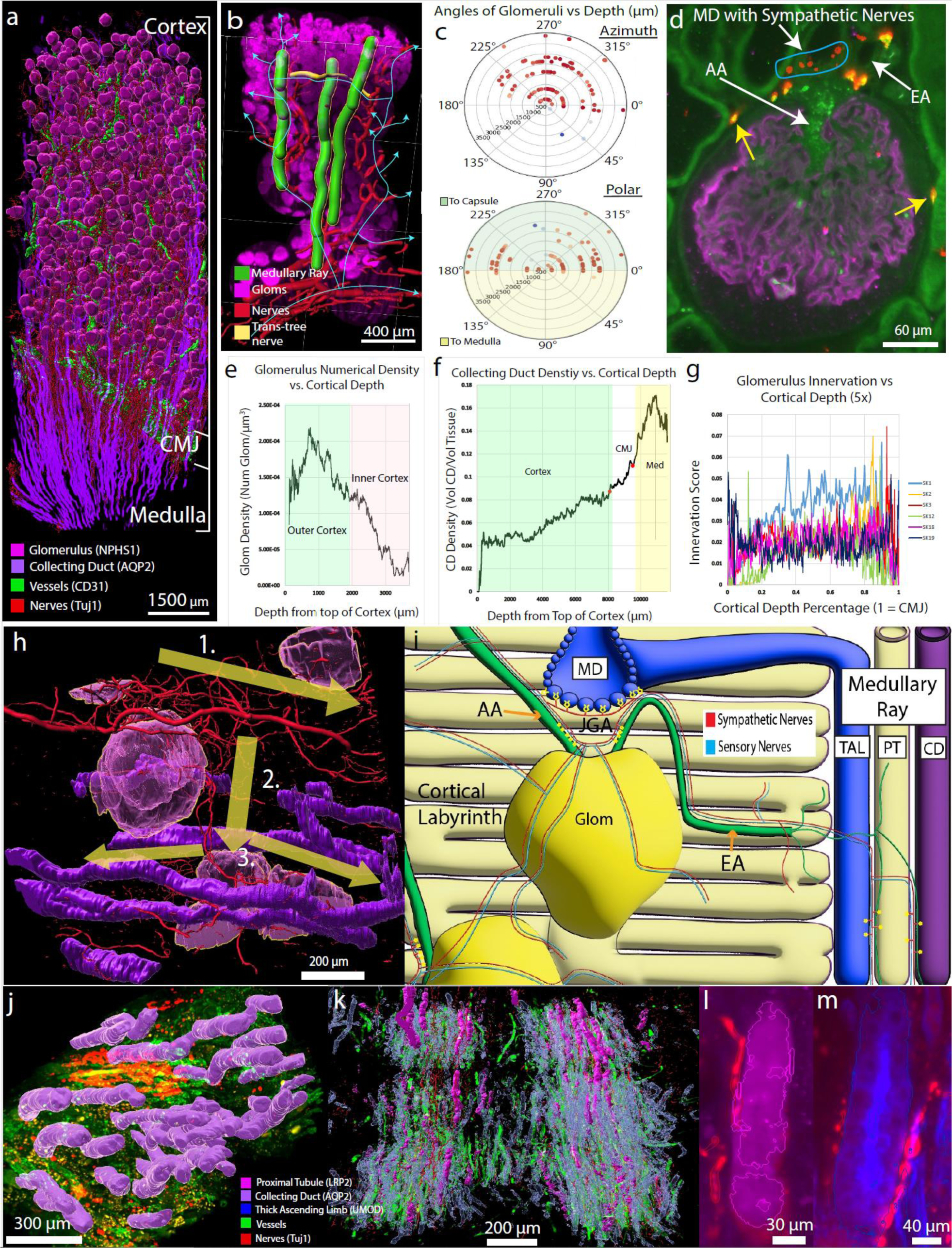
Three-dimensional morphometric analysis by LSFM. A) 3D LSFM image depicting organization of key structures along the cortico-medullary axis. The representative segmented image from a grossly normal 46 y/o male (Sample SK3) shows overall organization of glomeruli (magenta spheres) in the cortex, vessels (green) and nerves (red filaments) entering at the cortico-medullary junction (CMJ) from where they extend as branches in to the cortex or medulla with the nerves following the vessels till the glomerular vascular pole or along the vasa recta; the collecting ducts (purple) extend from outer cortex to medullary tip (see other panels and **supplementary movie 1**). B**) LSFM on neonatal pediatric kidney samples.** A segmented region of a 5x image depicting glomeruli (magenta), nerves (red and yellow), and medullary rays (green) demonstrated innervation patterns at a macroscale in the cortex spanning several lobules depicts that separate branches of nerves following the vessels into glomeruli further reconnect through glomeruli at various depths in the cortex (shown by line segments with arrows). Two distinct neural-trees can be observed branching upwards into the cortex from arcuate-artery associated bundles. C**) Radial heatmaps comparing the angle of the glomerular vascular pole (degrees) to its depth in the cortex (µm) using spherical coordinates.** Along the azimuth plane, glomeruli display a gaussian, two-quadrant bias in their orientations irrespective of depth (n=96 glomeruli). Along the polar plane, glomeruli display a bimodal, two-quadrant bias in their orientations irrespective of depth that demonstrates they tend to orient their vascular poles towards the kidney capsule, receiving vascularization from above (**Supplemental table S3** for source data). D) **Sensory and sympathetic innervation at the JGA and around glomeruli.** Th**e** 20x image is from a focal plane of a 3D LSFM experiment shows TUJ1-positive nerve (red) without overlapping CGRP innervates the entire macula densa (MD) while CGRP-positive sensory nerves (green) innervate only the flanking MD cells (also see **supplementary movie 3**). **E-G) Morphometry of FTUs.** The graphs demonstrate changes in morphological patterns of glomerular density (B, n=11; Supplemental Table S4), collecting duct density (C, n=8; Supplemental Table S5) and nerve density around glomeruli (D, n=6; Supplemental Table S6) from outer cortex towards medulla obtained from 5X LSFM images. Briefly, superficial glomeruli are more densely packed while corticomedullary glomeruli are arranged in separate columns around converging medullary rays. The collecting duct density gradually increases from superficial cortex towards CMJ where it rapidly rises and continues exponential rise till deep medulla consistent with convergence of collecting ducts from different lobules. Interestingly, nerve density around each cortical level of glomeruli showed an overall steady innervation density after an initial rise in the superficial cortex and a fall at the CMJ. This finding was further supported by manual and automated glomerular-nerve measurements in 20x volumes. **H) Relationship of nerves with glomeruli, juxtaglomerular apparatus and medullary rays.** The 20x volume of segmented glomeruli (magenta), collecting duct (purple), and TH-labelled sympathetic nerves (red) demonstrates interrelationship between these structures, the numbers (1-3) denote an example of nerve path. Here the nerve travels along the interlobular vessel (1) then projects along the afferent arteriole to vascular poles of glomeruli (2) from where this same nerve follows the efferent arteriole and projects to a nearby medullary ray containing collecting duct (3). **I. Summary illustration showing connectivity of nerves with various FTUs.** Both sympathetic (red) and sensory (blue) nerves travel with the AA to the JGA, where sympathetic nerves alone interact with the MD, but both types of nerve demonstrate bifurcation along the outer bowman’s capsule followed by further projections to other glomeruli, into the cortical labyrinth, back to the interlobular-vessel associated nerve bundle, or into the medullary ray where they most often innervate the proximal tubule and thick ascending limb, with less innervation along the collecting duct. Neural projections into the medullary ray appear to consistently occur beyond glomerular wrapping, but some projections into the cortical labyrinth appear independent of glomerular wrapping. Note that manual tracing also showed that the same nerve from the AA jumps to the efferent arteriole, travels along it while sending projections to the TAL of the same nephron, in addition to proximal tubules, and CD in the medullary ray (**supplemental movie 3**). **J-M) Segmented images showing neurovascular relationships with FTUs in the medulla (**s**upplemental movie 4).** J, The nerves are associated more richly with the vasa recta than collecting duct bundles. **K**, Unlike the CD, PT and TAL travel amidst the vasa-recta/nerve columns, receiving more prominent innervation. **L**, Innervation of a PT from a slice in K. **M**, A slice from the image in K depicting the innervation of a TAL.

As expected, the arcuate vessels entered the cortico-medullary junction (CMJ) and then projected interlobular arteries into the cortex that branched laterally into afferent arterioles (AA) in a step-wise manner from CMJ to outer cortex and ending at the vascular pole of each glomerulus and often branched to connect vascular poles of several closely associated glomeruli (**Figure 2a, supplementary movie 2**). The nerves also followed these AA that connected several closely located glomerular communities and common intralobular artery consistent with recent observations ^13^. The AA entered the glomerulus as glomerular capillaries and exited as efferent arterioles (EA) that further branched into capillaries surrounding the tubules or transitioned into the vasa recta into the medulla. The nerve trunks consisting of both sensory and sympathetic nerves follow this vascular pattern to the vascular pole, consistent with previous reports of the vascular pole as a heavily innervated site. However, we note that the distinct neural trees emerge from the arcuate vessels tracking along the interlobular vessels form horizontal connections between adjacent neural trees along rows of glomeruli laterally interspersed between medullary rays (**Figure 2b)**, suggesting secondary organization of neural activity on a macroscale. We next examined the anatomy, structure and cell type relationships of the vascular pole of the glomeruli as it harbors juxtaglomerular apparatus (JGA) ^14^, an important FTU in the kidney consisting of terminal AA, EA, extramesangial glomerular cells and MD; MD apposes the AA-EA with intevening extraglomerular mesangium ^15^. The JGA regulates blood flow to the glomerulus and hence filtration and operationalizes highly synchronized regulation of tubuloglomerular feedback, RAAS and positive and negative feedback between the sensory and sympathetic nerves ^16^. We first examined if the relationship of the topology of arteriole entry into the glomerulus was random or instead followed a specific orientation that fixed the JGA in particular orientation. We developed an algorithm to measure spherical coordinates of angular orientations of glomeruli with respect to their vascular poles from outer cortex to the CMJ in azimuth and polar axis. We found a two-quadrant, Gaussian directional bias in the azimuth axis, and a two-quadrant, bimodal directional bias in the polar axis where vascular poles were usually oriented “upwards” towards the kidney capsule at shallow angles throughout the cortex (**Figure 2c, Supplemental Table S3**). The bimodal polar orientations demonstrated that glomeruli finish maturity in one of two general orientations that may be hypothetically explained by them forming on opposite sides of a ureteric bud, resulting in two sets of glomeruli that point in opposite directions ^17^. Likewise, the Gaussian azimuth bias for the third and fourth quadrants may be a result of directionally-biased vascular development in the kidney. Since vessel structures arising from the renal artery must enter kidney regions from the same initial direction ^18^, glomeruli may be oriented towards these vascular inputs, inducing an angular bias.

We next examined 3D relationship of nerves with the MD and glomerulus. The nerves upon reaching the terminal end of the AA at the vascular pole bifurcated into two or more fibers none of which follow the AA into the glomerulus itself. In addition to connections with the vasculature, the nerves pass beneath the MD at the JGA where Synapsin I staining revealed a high-density of synaptic varicosities indicating contact points with the MD **(Supplemental Figure S1j)**. Another branch of these fibers moved on from the vascular pole and travelled avascularly along the outer layer of the Bowman’s capsule (BC) surface, wrapping the glomerulus like a net, consistent with recent observations ^13^, before moving onto additional targets (see later) and avoiding the urinary pole **(Supplementary movie 2).** While both sensory (CGRP+) and sympathetic (TH+) nerves were present at the vascular pole, we noted a distinct pattern of MD innervation by each **(Figure 2d, Supplementary movie 2, z-axis 2D views).** The TH+ fibers innervated all MD cells spanning the MD while CGRP+ nerves only targeted the flanking MD cells of the MD region indicating a novel basis for some of the sensory-sympathetic feedback mechanisms in response to alterations in tubular function and blood flow and regulation of the renin angiotensin aldosterone system (RAAS) ^15,19^.

Quantitative analysis revealed a decrease in glomerular density from outer cortex to corticomedullary junction where converging medullary rays occupied more space ^20^ (**Figure 2e, Supplemental Table S4**).

Collecting ducts (CD) converging radially gradually occupied greater density **(Figure 2f, Supplemental Table S5**) before spiking in density upon reaching the corticomedullary junction (CMJ). This resulted an overall broccoli-like glomerular distribution, with distinct glomerular columns in the juxtamedullary cortex where converging rays took more space, but dense clumps in the superficial cortex (**Supplemental Figure S1a**). This glomerular pattern in the adult kidney is consistent with a rapid increase in nephron number later in human kidney development where they grow as arcades emanating from the branching ureteric buds ^21^. The CD density continues to rise throughout medulla consistent with merging of CDs from multiple medullary rays and renal lobes. In contrast, the nerve densities at different depths of glomeruli did not vary outside of an initial rise in the very superficial cortex (**Figure 2g, Supplemental Table S6).** 20x innervation quantification across cortical depths likewise supported the idea that glomerular innervation was largely invariable across cortical depths **(Supplemental Figure S1d, Supplemental Tables S7, S8)**, implicating that the slight rise observed in 5x superficial cortex was a factor of diminishing signal around thinner nerves rather than a true anatomical feature.

### Nerves connect FTUs of the same nephron and others in the community

We next traced the path of the nerves after their tracking along the AA and innervation of JGA components and the glomerulus **(Figure 2h, Supplemental movie 3).** Groups of closely apposed glomeruli often demonstrated several nerve connections to one another after leaving the outer layer of the Bowmans capsule (BC), serving as communities in a neuronephron-network. Concerning tubules, branches at the JGA jumped from the AA to the EA, before moving beyond the glomerulus to the medullary ray and innervating other nephron segments including the PT, CD, and TAL for some distance but not reaching the CMJ. In the medullary ray, the same nerve branched to innervate adjacent TAL, PT, and CD. Although innervation of TAL was confirmed to be of the same nephron, we could not conclude this about the PT or CD due to their long paths exiting the FOV, however there was some suggestion of this due to the nearby adjacency of the innervated PT and the PCT of the nephron in question, and the connecting tubule of the innervated CD being oriented towards the DCT but exiting the volume before connecting. These data demonstrate that the same nerve connected the glomerular periphery and tubular segments from the same nephron, setting novel precedence for close communication between these structures **(Figure 2i).** We also noted that there was cross nephron tubular segment connectivity by nerves that are a part of glomerular communities described above indicating quite a sophisticated system of cross talk among these FTUs **(Supplemental Movie 3).** Within the ray, PTs and TALs appeared the most prominent targets for innervation, often receiving parallel-nerve branches, while the CD appeared innervated only at specific points. More nerve targets downstream of glomeruli included short-lasting projections into the cortical labyrinth, including PCTs innervated by branches from its glomeruli and other glomeruli closely linked in the neuroglomerular network, and by nerves that returned to larger nerve bundles moving with interlobular vessels. While glomeruli and their associated nephron structures were the primary target of nerves in the cortex, the nerve bundles also sent numerous short-lasting projections into nearby cortical labyrinth, unassociated with glomeruli, as they progressed through the tissue (supported by Synapsin I and TUBB3 staining). Synapsin I in particular (**Supplemental Figure S1I**) emphasized previous studies that showed nerves in the kidney cortex utilized consistent en-passant neuroeffector junctions ^22^ together suggesting that nerves in the kidney cortex provided some degree of innervation at all points traversed.

### Innervation pattern in the medulla

In the medulla, vasa recta projected downward from arcuate vessels in interspersed columns, followed by networks of nerves projecting from the arcuate-associated nerve bundle. Organization of nephron structures within the medulla differed from the medullary rays, with both the proximal tubule and loop of Henle travelling amidst vasa recta-nerve columns, while CDs moved apart from the columns in separate, largely anervous bundles of the long loop nephrons from deep cortex (**Figure 2j, k**). Consequently, innervation in the medulla appeared to primarily target the vasa recta, PTs (**Fig 2l**) and loop of Henle (**Fig 2m**) far more than the CD **(Supplemental Movie 4)**.

### Unique patterns of neuronephron connectivity

The extensive innervation revealed above prompted us to further examine connectivity between nerves and glomerular communities to glean insights into how kidney FTUs may work in a coordinated manner. We use the concept of a network as a collection of objects, known as nodes, and their connections to one another, known as edges. Within networks, cohesive groups called ‘communities’ can be formed, which relate to a system’s functional modules ^23^. Groups of nodes organized into communities may together preform distinct, semi-autonomous functions within the wider network ^24^.

To discover neuro-glomerular networks from 3D LSFM imaging data we developed a novel algorithm where we treated glomeruli as nodes and their neural connections as edges using both manual tracings and automated network detection after the nerves have left the vascular pole **(Figures 1, 3, see methods).** We discovered repeating patterns or motifs of glomerular innervation within and across communities (**Figure 3**). Network motifs represent network patterns that repeat more frequently than in a random network, which in biological systems are often associated with discrete information processing substructures of a network ^25^. We identified seven motifs of neuronephro-networks that we broadly group into intra-community motifs describing basic glom-nerve relationships **(Figure 3a-c),** community motifs describing patterns of community aggregation **(Figure 3d, e),** and inter-community motifs, which describe the relationships between communities **(Figure 3f, g**); illustrated in **Figure 3h** and **Supplemental Movie 5**.

**Figure 3.**
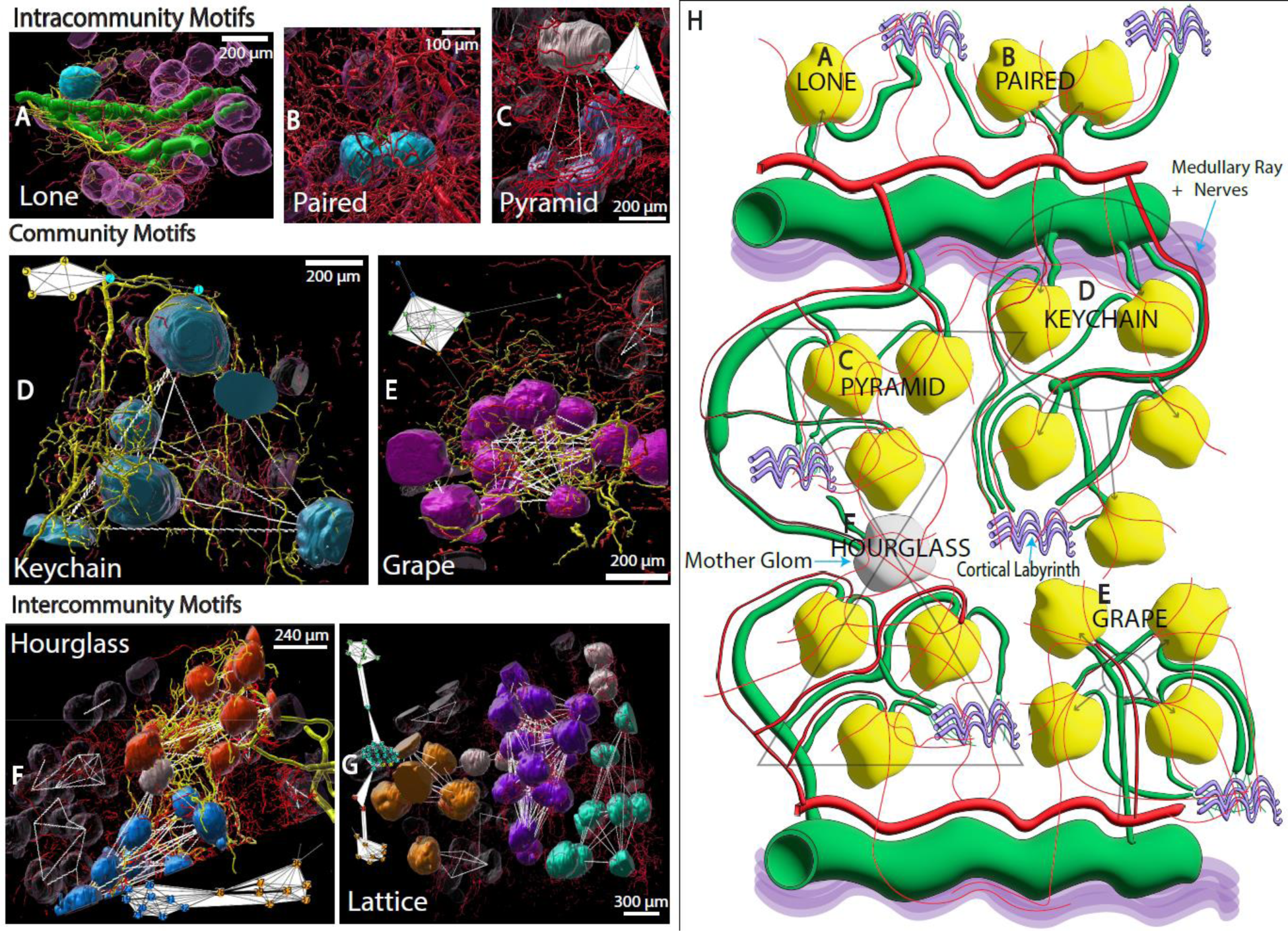
**Neuro-glomerular network motifs. A-C, M**otifs existing within a community, **D-E** Community motifs, and **F-H** represent community relationships. **A)** A lone glomerulus (blue) receives nerves (yellow) directly projecting from an interlobular vessel (green), without direct neural projections to other glomeruli. Nonparticipating nerves and gloms are shown in red and magenta, respectively. **B)** Paired glomeruli (blue) are closely apposed and are vascularized by an afferent arteriole (green) that forks into two vessels immediately before entering both. They share a tightly linked neural network along their vascular poles and wrapping the outer layer of the bowman’s capsules. **C)** A pyramid motif occurs when one glomeruli (white) appears to be upstream in a network to several other glomeruli (blue). The network graph of the same glomeruli is overlayed atop this, and all following, motif images. **D)** A keychain motif (blue) occurs in low glom density tissue when a branching neural projection from the interlobular vessel spreads between several glomeruli before looping back to its nerve bundle in a circle. Keychain motif appeared as typically five to seven nodes (representing glomeruli) of equivalent degree linked in a ring **E)** A grape motif (solid magenta) occurs in high-glom density tissue when glomeruli oriented around an interlobular vessel surround a centralized nervous hub, whose projections link multiple glomeruli (8 or more). **F)** An hourglass motif (solid-colored glomeruli) occurs when two distinct neuro-glomerular communities which receive distinct vascular inputs are unified by one or a few vertex glomeruli that sit between communities, referred to as mother glomeruli. **G)** Lattices (solid-colored glomeruli) occur when multiple communities are unified by mother glomeruli. **H)** A cartoon depicting several of the neuro-glomerular motifs and their relationships with other nephron structures. Beyond joining glomeruli into networks, nerves may establish additional connections between nephron structures in the cortical labyrinth or medullary ray the letters correspond to the images A-F. See supplemental movie 5.

We were particularly intrigued by the inter-community motifs. The hourglass (**Figure 3h, Supplemental Movie 5—0:36-1:20)** structure was observed when two grape or keychain communities had their nerve networks joined by one or more central glomeruli. Significantly, each community was observed to have a distinct interlobular blood vessel source and community-specific projections into the medullary ray; the central glomeruli had additional projections into the medullary ray from both communities in the hourglass motif. Consequently, these nerve networks and central glomeruli joined together otherwise discrete nephrovascular communities, possibly synchronizing activity between adjacent glomerular communities. In the network, hourglasses appeared as two dense groups of high degree nodes joined by one or a few nodes that existed between these communities. Attached to the hourglass, additional interglomerular connections could also be seen to join lower-degree nodes that themselves existed as intra-community motifs such as pyramids.

Furthermore, analysis of the large 3D volumes revealed the lattice motif when three or more communities joined together by central glomeruli **(Figure 3g, Supplemental Movie 5—2:24-2:50).** This structure represented the largest macroscale pattern we could observe in these networks. Lattices were not restricted to linear connections between multiple communities; for example, 3 communities could be joined to one another in a triangle structure. In dense 5x networks, this often resembled a classic spoke-and-hub network structure. Organization like this has been described in real world systems where interconnected networks enable information or material flow with processing at central critical nodes (hubs) ^26^. This organization is advantageous as it simplifies networks of scale by optimizing hubs, a trait relevant to kidneys that must control up to a million FTUs simultaneously, with a notable weakness being a vulnerability to the disruption of hub nodes. This may suggest the loss of these central glomeruli to be more deleterious to neuronephro-networks as compared to other glomeruli. Given that the community-unifying pathways transverse along central glomeruli, we hypothesize that these nodes may be of greater importance to network function and subsequently define these objects as *‘mother glomeruli*’. A summary of all these motifs and their relationships with other nephron structures with the mother glomeruli at the center of an hourglass is illustrated **(Figure 3h).**

### Characterization of neuro-glomerular networks across scales

We next performed a number of analyses to determine the robustness of the neuro-glomerular network across scales at both 5x and 20x resolutions, pattern at different glomerular densities and types of nerves **(Figure 4).** To reveal networks across communities at low power, the main nerve trunk to which all structures are connected were removed. One way to describe communities within a network is through its “modularity”. A modularity closer to 1 is indicative of more statistically ‘surprising’ community structure, with greater intracommunity as opposed to intercommunity connectivity, while a value closer to 0 represents a more random arrangement ^24^. Higher-glomerular density samples with abundant nerves exhibited a single, large and highly modular interconnected component, demonstrating an abundance of secondary interglomerular nerve connections beyond the trunk. This component often contained the majority of the glomeruli that were previously present if the trunk was not removed **(Supplemental Figure S2b).** Lower-density samples with more diffuse nerves often had many glomeruli released into small components or removed from the network entirely if the trunk was removed, but left a few medium-sized components representing modular hub-and-spoke entities **(Figure 4a, Supplemental Figure S2c).** It subsequently appeared that samples did vary in their community synchronization, with low- glomerulus/nerve density tissue typically possessing a greater degree of more-independent modules while denser cortices had more abundant interconnections. Nonetheless, hub-and-spokes dominated amongst any larger components that remained in all trunkless networks.

**Figure 4.**
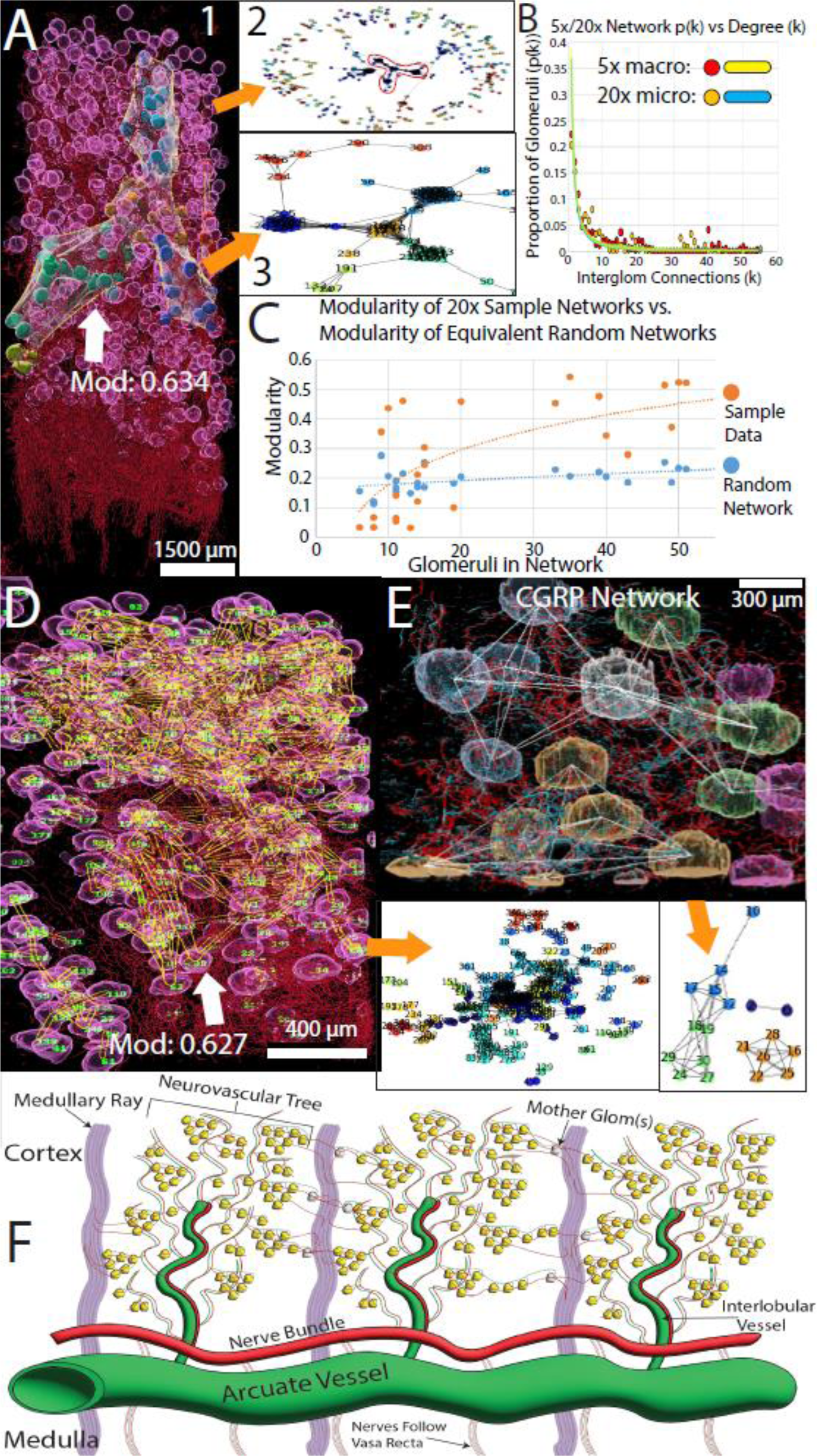
**Characterization of neuro-glomerular networks at both 5x macro and 20x micro scales**. **A1-3) Network analysis at 5X magnification reveals highly modular neuro-glomerular connectivity. A1**, A representative 5x segmented image of a reference sample (SK3, PPID 3778) containing spheroid glomeruli and red filament nerve segmentations (TUJ1 labeled) spanning the cortex; the largest interconnected network component is highlighted with the glomerular communities colored solid and the arrow indicating its modularity (0-1 strength of community structure) score. **A2**, Connectivity map of A1 with each glomerulus numbered and the same large component highlighted in A1 is circled in red. **A3**. Enlarged view of the connectivity map of the highlighted component. The community colors correspond to the colored communities in the image in A1. **B. A graph depicting the proportion of glomeruli (p(k)) in a network that contain ‘k’ neural connections to other glomeruli at both 5x and 20x resolutions (**sample SK1, PPID 3785). The 20x distribution (n = 222 glomeruli) is shown in orange and its blue trendline (R^2^=0.800) has the equation p(k) = 0.3664x^-1.418^ while the 5x distribution (n = 440 glomeruli) is shown in red and its yellow trendline (R^2^=0.852) has the equation p(k) = 0.3833x^-1.402^. These networks demonstrate scale-free properties by both satisfying the baseline assumption that the exponent α > 1, and a near-identical degree distribution at micro- and-macro scales. **C)** A graph comparing the modularity of reference 20x neuroglomerular networks in this study to the number of glomeruli contained in said 20x network in orange, while also comparing the modularities of random networks of equivalent size in blue. At <10 glomeruli, networks were observed to be low modularity due to often lying within the bounds of a single community, however modularity rapidly spiked between 10-20 glomeruli and continued to rise slowly, as more communities were involved. The largest 20x network contained 222 glomeruli and has a modularity of 0.627. The reference networks are less modular than random when within communities, but more modular when encompassing multiple communities, demonstrating that glomeruli are controlled by a modular, non-random network structure. **D) Neuroglomerular connectivity map at 20X magnification.** Representative 20X image from same sample in the graph in B from FOV #8 with segmentation depicting glomeruli (magenta), nerves (red), and network connections (yellow) encompassing 370 glomeruli. Trunkless network analysis yielded 222 glomeruli connected in a single, highly- modular network. 2. **D’** The corresponding network graph for D. The network connectivity map with each glomerulus is numbered and trackable in 3D. **E)** CGRP-labeled sensory neurons contribute to the neuroglomerular connectivity map. Representative image from a single 20X FOV from the same patient as in D (sample SK19, PPID 3778) depicts segmented glomeruli (spheroids) arranged in a network using CGRP-labelled sensory nerves (blue filaments). The large component shows a pattern of two glomerular pyramids joined by mother glomeruli in an hourglass motif. Tuj1-labelled nerves for TUBB3 are shown in red. **F)** A cartoon depicting macro interactions between neurovascularization, glomeruli, and the medullary ray. Neural trees follow interlobular vessels from the arcuate vessel into the cortex to innervate glomeruli. They form glomerular neuro-communities that can be unified via mother glomeruli across the cortex and different lobules to as connected communities downstream of separate neurovascular trees. These connections often occur along glomeruli sitting along lateral pathways between medullary rays, which themselves contain parallel-oriented projections downstream of glomerular innervation.

Within 20x FOVs, similar patterns emerged, however the modularity and the strength of community structure varied proportional to the number of glomeruli captured in the image and these observations show a non-random distribution **(Figures 4b-d, Supplementary Table S9).** This modularity trend was the result of images with a few glomeruli typically containing only small intracommunity/community level structures, while 20x images with many glomeruli captured multiple communities in one FOV. Likewise, a sharp increase in modularity can be observed between 20x images containing between 10 and 20 glomeruli and also supported in comparative analysis in random networks of equivalent node/edge count. Thus, below 20 glomeruli, samples demonstrated inconsistent community structure that could be less-than-random, but that images with >20 glomeruli samples yielded networks with greater-than-expected community organization. These data therefore suggested that community-level motifs (keychain and grape) contained between 10-20 glomeruli in general, which together formed intercommunity structures (hourglass and lattice) with seemingly systemic rather than a random community organization (Welch’s t-test p = 0.013). The intercommunity connectivity diminished as glomeruli became less dense, but these patterns could nonetheless be observed in both dense and diffuse glomeruli tissue so long as sufficient number of glomeruli were incorporated in the image. Nonetheless, these samples differed in their apparent network involvement, with communities linked at more frequent points in the high-density sample, implicating multiple mother glomeruli, but the low-density sample demonstrating sparser interlinkages containing only one or two mother glomeruli.

The observation that the trendlines of glomeruli number and networks at 5X and 20X resolutions from same sample overlap suggest that these are scale-free networks **(Figure 4b).** A scale-free network is one not defined by its size or scale, but rather the existence of highly important nodes or hubs ^27^, consistent with the spoke and hub structure visible in our networks. A scale free network generally has the property of the fraction of nodes with degree k following a power law distribution p(k) = k^-(α), where α>1, and 3> α >2 being more strongly scale-free ^28^. Of the reference 5x networks analyzed for this property, seven of nine had degree distributions that satisfied the baseline assumption of α>1, with all trendlines having r^2>=0.8, which also applied to our largest 20x network (**Figure 4d, Supplementary Table S10**). It should be noted that only one network satisfied the stronger assumption of 3 > α > 2. Such scale-free, hub-oriented networks carry the important advantage of being strongly resistant to network damage due to accidental or random failures, as the loss of non-critical nodes will scarcely impact scale-free network function as compared to a random network.

However, such networks display a critical weakness as the loss of even 1.5 to 5% of critical nodes can splinter a network into disconnected subgroups. For example, this can be seen in cell metabolic pathways, where mutations in the majority of proteins may hardly affect a cell, but the knockout of the few key metabolic proteins will have a higher chance of causing cell death. ^27^. On the other hand, a heavy-tailed, inverse power distribution, where a large proportion of nodes were low-degree is suggestive of a complex network more prone to extreme behaviors, such as complete-network activation, when compared to a random arrangement ^29^, a factor that would aid any simultaneous management of parallel nephrons.

Finally, we examined if sensory nerves (CGRP+) contribute to these communities as it would support sensory-sympathetic neural cross-talk across glomeruli and communities in the kidney to modulate neurophysiology of kidney that has largely been implicated to the vascular pole at the JGA and ^30^. Manual nerve tracing in these 3D volumes and CGRP network results demonstrate CGRP-stained sensory nerves themselves are sufficient to generate a neuroglomerular network with some level of community structure (**Figure 4e**), consistent with our observations of CGRP nerves wrapping and extending further between gloms; TH+ nerves also contributed to the community network, appearing to almost never diverge from Tuj1 stain unlike CGRP **(Supplemental Figure S1g),** suggesting a greater abundance of sympathetic nerves overall. A macro-scale drawing summarizing the above neuronephro-networks utilizing mother glomeruli is shown in **Figure 4f**.

### Neuro-nephron patterning across postnatal life span

We next investigated changes in neurovascular nephron organization of the human kidney over life span, including newborn to aged kidneys (1d neonate, 1mo infant, young adult 18-40yo, adult 40-75 yo and aged >75 yo), using LSFM to understand how structural changes may impact kidney function **(Figure 5).** One advantage of pediatric kidney tissue due to its small size was that it enabled visualization of large areas spanning the whole kidney lobe by LSFM. In the neonate kidney, glomeruli were densely packed, showing a well-organized cortex and a pattern reminiscent of radial nephron development during branching morphogenesis ^31^ **(Figure 5a; Supplemental movie 6).** In the cortico- medullary axis, glomerular columns were interspersed in columns between medullary rays that merged into clumps as the medullary rays blended in deeper cortex. This is consistent with glomerular arcades forming in deep cortex, where a single ureteric bud branch induces multiple nephrons to rapidly increase nephron number ^21^. Interestingly the neonatal glomeruli were bean-shaped compared to spheroidal in the adults (**Figure 5b)** and their polar angles lacked the apparent bimodality of the young adult (neonates fail to reject Kolmoglorov- Smirnov normality test; p = 0.08 vs. young adults reject K-S normality test; p =1.8e-4), possibly explained by the compact packing of neonatal glomeruli that later settle into shallower angles by radial enlargement of the kidney **(Figure 5c).** While the vascular networks in the cortex and medulla was well established, the nerve network was visually and quantitively immature relative to the adult kidney. At 1d, the nerve bundles followed the arcuate artery and branched into the cortex alongside interlobular vessels, but had not yet developed connections to glomeruli or other nephron structures **(Supplemental Movie 7)**. In the medulla, nerves followed vasa recta distal to collecting ducts and did appear to interact with the PT/TAL but with less overall complexity than the adult tissue. The medulla was visually more compact with structures in the outer and inner medulla less defined ^32,33^. **(Supplemental Movie 4)**

**Figure 5.**
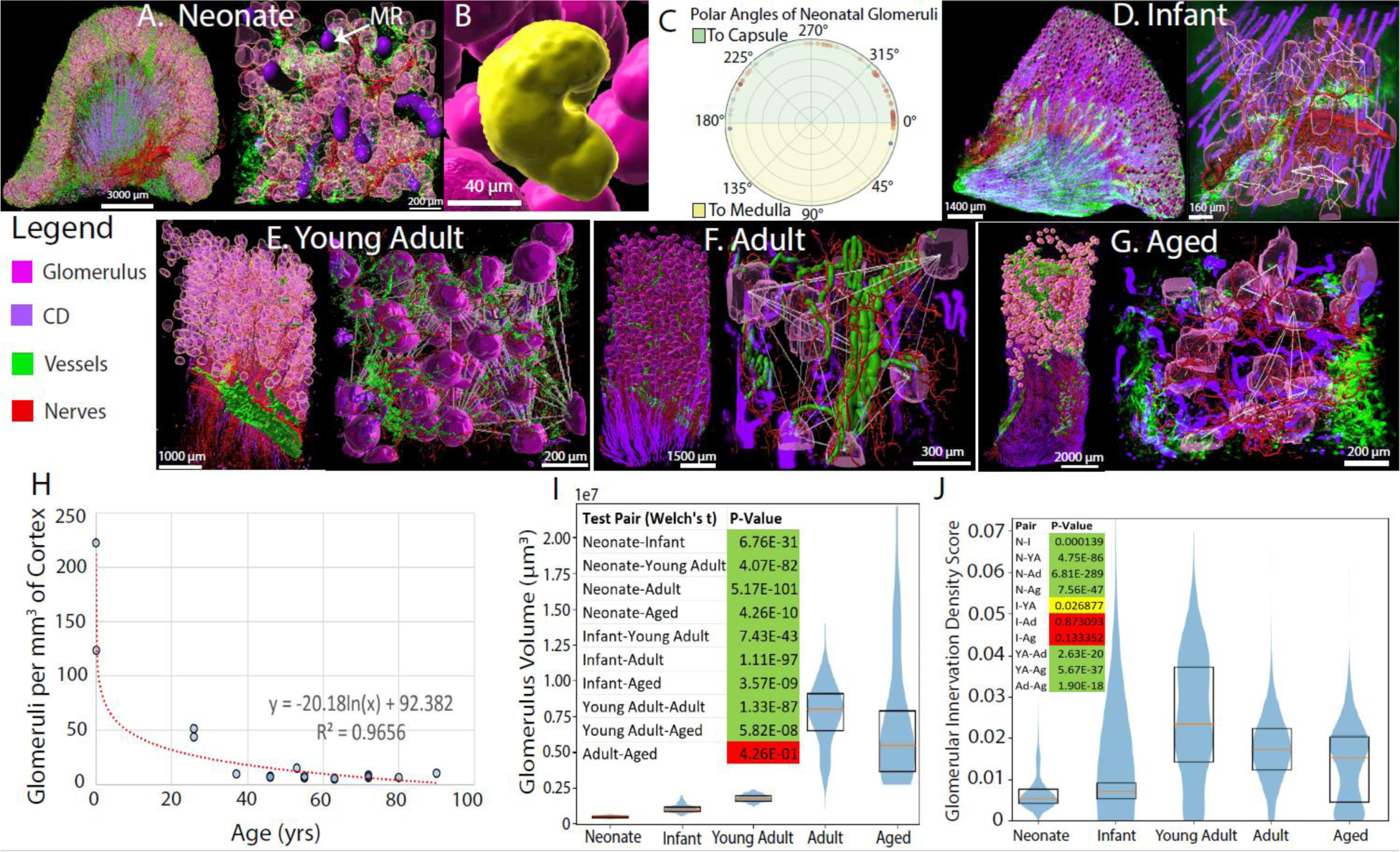
3D Neuroglomerular morphometry across life span. A) Neurovascular and nephron anatomy in a neonatal kidney. The 5X image of an entire neonatal renal lobe (left, sample SK14, PPID-3961; 1d) and corresponding 20x image of the cortex(right) shows that the nerves follow the vessels until afferent arteriole where their final connectivity to the vascular pole is underdeveloped and projections to the medulla aren’t established. Glomeruli and medullary rays (MR - shown in purple) are packed tightly. **B)** N**eonatal glomeruli are bean-shaped rather than spherical. C) Polar angles of the vascular pole in the neonate are not bimodal. D) Neuro-nephron connectivity is beginning to appear in an infant kidney** (Sample SK16, PPID3730, 1m). Illustrated is a 5x image of an entire renal lobe (5X, left) and cortex (20X, right) showing neural connections to and between glomeruli are more established including neuroglomerular networks (connections between glomeruli - white lines). Glomeruli show increase in size, less dense and are spherical. **E) Neuro-nephron connectivity in a young adult kidney at 5X (Left) and 20X (Right) (S**ample SK2 from PPID3785, young adult**) shows abundant modular network connections**. Note that the glomerular density has decreased substantially compared to infant, while volume has increased. **F) Neuroglomerular connectivity and glomerular density decreases in adult kidneys (Sample SK3, PPID3778, 46yo).** (Left) A 5x image of renal lobe. (Right) A 20x image of adult cortex. **G. Aged adult kidney shows a range of glomerular size from atrophic to hypertrophic, increased glomerulosclerosis and decreased neuronephron connectivity** (Sample SK12, PPID3900; 75+). (Left) A 5x image of cortico-medullary renal slice and (Right) a 20x image of aged adult cortex also shows that the glomerular density is comparable to adult samples. **H. Glomerular density rapidly decreases from neonatal years to young adulthood and then gradually across adult life span.** circles - individual sample. **I. Violin plots depicting glomerular volumes measured at 20x resolution for the different age groups.** Neonates, infants, and young adults depict tight distributions that show steadily increasing in volume with age. Adults demonstrate a mostly a gaussian distribution, however, aged adults show a distribution with mostly declining volume by tuft shrinking, except for a right skew caused by hypertrophic glomeruli. A Welch‘s t-test found highly significant differences (p < 0.01) between all groups except between the adult and the aged adults. **J. Violin plots depicting glomerular neuroinnervation sampled across different regions of cortex at 5x resolution.** Neonates demonstrated little innervation amongst all glomeruli. In Infants, neural connections to glomeruli have begun to form, resulting in a right-skew. Neuroinnervation peaked in young adults before gradual decline into aged adult. The aged sample demonstrates prominent bimodality, with the majority of glomeruli resembling the adult innervation distribution, but with a low-peak caused by low-innervation glomeruli that have begun to develop. A Welch’s t-test found highly significant differences (p < 0.01) between all groups except when comparing infants with young adults, which demonstrated standard significance (0.01< p < 0.05), and when comparing infants to adults and infants to aged adults, which demonstrated no significance.

As kidney development progressed, the glomerular density decreased, volume increased steadily till young adult hood before exponentially increasing in the adult age group **(Supplemental Table S11).** In the aged kidneys, there was increased as well as decreased glomerular volume observed likely due to hypertrophy of resident glomeruli to compensate for progressive nephron loss supported by previous studies ^34^ **(Supplemental Table S12).** Importantly, the neuro-glomerular networks and community connectivity were noted at 1mo and are well-established into connected communities and mother glomeruli in the young adult **(Figures 5 d-g, Supplemental Figure S2b).** Interestingly, neuroglomerular networks appear more diffuse in adult kidneys, with far lower secondary glomerular connectivity beyond the nerve trunk **(Supplemental Figure S2c, Supplemental Table S13)**. Large modular structures could be detected but were demonstrably smaller in their glomerular inclusivity compared to similar structures in the young adult. Patterns of innervation and organization were otherwise comparable. Adult glomeruli displayed some signs of early obsolescent sclerosis, where the tuft peels away from the tubular pole ^35^. In aged individuals there was a notable heterogeneity in glomerular shapes including small-shrunken sclerotic and oblong that may explain the increased density observed in this age group ^35^ **(Supplemental Figure S1f)**. In addition, the aged group demonstrated a 5x nerve network of high modularity and comparable glomerular incorporation to the young adult **(Supplemental Figure S3)**, as opposed to the more diffuse adult networks, despite having glomerular density comparable to adults. These observations are supported by quantitative analysis of trends across life span in glomerular density, volume and innervation (**Figure 5h-j, Supplemental Movie 7).**

### Disease states affect neuro-nephron networks

We next investigated if neuroglomerular networks are altered in patients with kidney disease or pathology that may provide new insights into structural basis for associated pathophysiology. We examined neuroglomerular networks in sclerotic glomeruli in aged kidneys, glomeruli in hydronephrosis near and away from the hydronephrotic area, and in a patient with diabetes. We discovered three types of alterations in neuroglomerular networks that may be maladaptive **(Figure 6).**

**Figure 6.**
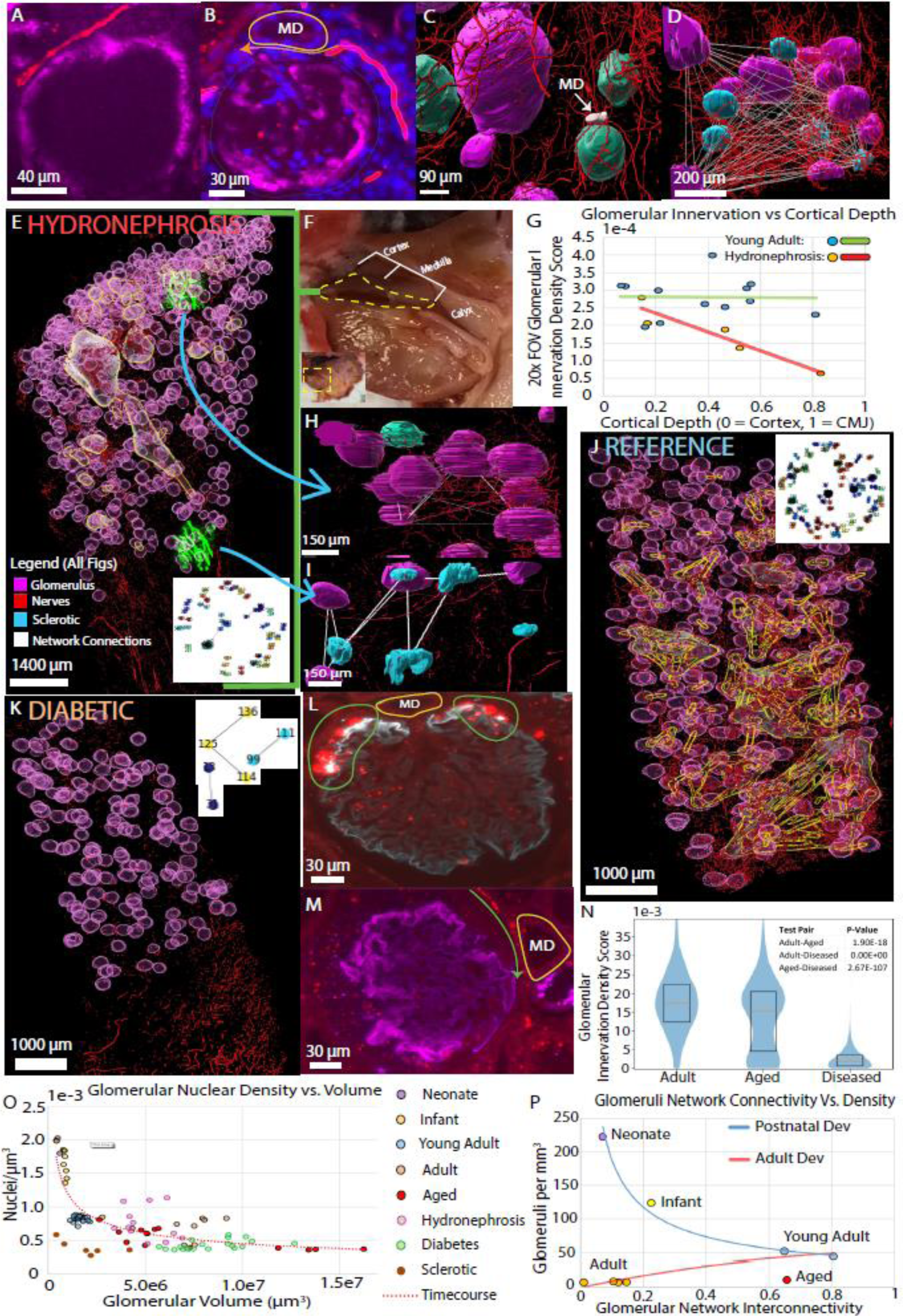
A**ltered neuroglomerular networks and morphology with age and disease using light sheet fluorescence microscopy**. **A-B) Periglomerular and vascular pole innervation persists in sclerotic glomeruli (>75yo).** Due to global glomerulosclerosis no nephrin staining (magenta) is seen except in the outer rim of bowman’s capsule (magenta) that is nonetheless receiving nerve fibers (red) in A. An example of sclerosed glomerulus with a few cells remaining still has innervation at the vascular pole. The path of the nerve around the macula densa (MD) has been drawn with an orange arrow; nuclei (DAPI stain blue). **C-D) 3D illustration of connectivity of sclerotic glomeruli in the community of nonsclerotic glomeruli.** Note size difference between sclerotic (cyan) and nonsclerotic glomerulus (magenta), the arrow points to the macula densa (MD) of the same glomerulus in B; all sclerotic glomeruli receive innervation and are in the community nerve network (white lines in D). **E-I) Neuroglomerular connectivity is affected more severely in regions closer to dilated collecting system in hydronephrotic kidney** (Sample SK8, PPID3909, 55yo). Segmented image of cortex and medulla (E) from a region (yellow dotted line) next to dilated calyx shown in the gross image in F depicts neural network connections by yellow/white bubbles alongside an overlayed network graph. The two volumes marked in green in E are shown at high power in H and I representing outer cortex and next to medullary junction, respectively, with visibly reduced neural connectivity compared to reference samples. Cyan - sclerotic glomeruli remain in the network. G. A scatterplot of glomerular innervation density shows in reference sample this remains constant along the C_M depth, whereas, in the hydronephrosis sample it decrease along cortical depth (n = 12 FOC from reference - 612 glomeruli; 5 FOV from hydronephrosis - 42 glomeruli). **J**. Adult reference sample (SK18; PPID3778, 46yo) with overlay of network graph demonstrates visibly greater network participation compared to diseased samples. **K-M) Neuroglomerular connectivity in diabetic kidney disease sample is markedly reduced** (Sample SK9, PPID3702, age 63). The network graph is overlayed.in K. L and M show examples of 2 different glomeruli with L showing an abnormal punctate pattern of nerve staining rather than filamentous appearing nerves at the vascular pole and M shows a fiber at its vascular pole passing beneath the MD (traced in yellow), demonstrating that complete neural disruption is not apparent for all glomeruli in this diabetic patient. **N) Violin plots of nerve density comparing adult, aged and diseased samples.** Reference adults have a Gaussian distribution while aged samples have a similar median but have acquired a bimodality caused by low-innervation glomeruli; diseased patient glomerular-innervation is significantly reduced (Welch’s t-test P values are indicated). **O) Summary of relationship between nuclei density and glomerular volume across life span and disease.** The scatterplot shows the nuclear density decreases in glomeruli with age as their volume increases likely due to age related cell loss and hypertrophy compensating progressive glomerular loss. The diabetic patient showed glomeruli that trended slightly larger and with lower cell-density on average compared to the adult group. The hydronephrotic patient, generally exhibit smaller glomeruli than the adult group but with mostly similar cell densities. Sclerotic glomeruli from the aged group (brown) had both low cell densities and diminished volumes. **P) Relationship of glomerular density and neuroglomerular connectivity across life span.** From neonate to young adult, the glomerular density decreases as the kidney grows but the network connectivity increases as nerves develop. However, in the adult and aged there is decrease in glomerular density, likely due to loss of glomeruli with age, decreased connectivity in less dense glomerular communities and a paradoxical increase in aged networks likely due to an overconnected network containing sclerosed glomeruli.

*The overconnected network.* Sclerosed glomeruli do not function but our aged patients surprisingly maintained substantial innervation of sclerotic glomeruli, incorporating them into their neuroglomerular networks in low density regions (**Figure 6a-d**). Even MD associated with the sclerotic glomeruli had nerve connectivity **(Figure 6 b, c).** Keeping nonfunctional glomeruli in the network could be maladaptive by feeding the apparently-intact network “bad” information and affect kidney homeostasis. One explanation for this pattern would be that as sclerotic glomeruli accumulate with age, the network responds with additional connectivity as an attempt to over-compensate and also not disrupt the highly connected network by loss of innervation to the sclerotic glomeruli.

*The regionally-disturbed network.* We examined neuroglomerular network in outer cortex and inner cortex closer to region of dilated calyx in a kidney with hydronephrosis in comparison to a reference kidney (**Figure 6e-j, Supplemental Table S7).** The hydronephrotic network was much reduced compared to the reference kidney and quantification of glomerular nerve density showed progressive decline from outer cortex to deeper areas closer to the hydronephrotic segment whereas the neuroglomerular density remained constant throughout the cortex in reference kidneys of similar density. There was increased number of sclerotic glomeruli in deeper cortex but they still retain connectivity to the glomerular network.

*The globally-disturbed network.* We also examined neuroglomerular networks in patients with diabetic kidney disease to gauge if there are innervation abnormalities in the kidney similar to neurodegeneration in neuropathy **(Figure 6k-m)**. Remarkably, small fibers were observed at the JGA but remaining areas displayed punctate pattern of Tuj1 (TUBB3) staining and no distinct fibers reminiscent of axonal neurodegeneration ^36^. To further increase confidence in reduced innervation in diabetic kidneys we performed confocal 2D immunofluorescence on 15 additional patients (5 reference, 5 diabetics with mild CKD with no kidney dysfunction and, 5 diabetic patients with severe CKD). The patients with severe CKD showed marked reduction in glomerular innervation compared to reference and mild CKD diabetes samples while the mild CKD patients showed an intermediate phenotype (**Supplemental Figure S3a, Supplemental Table S14)**. An acute condition such as AKI did not exhibit any reduction in neuroglomerular connectivity, suggesting that the reduction in network is correlated with chronic kidney conditions and thus may explain fluctuations in vital and metabolic profiles **(Supplemental Figure S3b).**

Manual and automated quantification of the above trends in disease and also across lifespan are depicted in **Figures 6 n and o and morphological variations in glomerular density (Supplemental Table S15) and networks in Figure 6p (Supplemental Table S13).**

## Discussion

3D multiplexed molecular imaging poses unique challenges in human solid organs due to complex architecture and matrix. Even if the methods of clearing and immunofluorescence are overcome, the amount of data generated pose significant challenge in analyses to understand structure-function relationships. The application of 3D LSFM to human kidney samples and analytical approaches presented here are key advances technologically and conceptually that are applicable to other organs. Our approach provides novel insights into how 3D structural organization permits various functions and explain kidney pathophysiology. In addition to confirming overall organization of the main neurovasculature branches and nephron components in previous anatomical studies, we provide novel findings on how these relationships and anatomical organization change from neonatal to aged time periods, such as the finding that neural connections to cortical nephrons develop postnatally. We further report neuronephron patterning in diseased samples or with abnormal pathology discovering several changes in cortico-medullary anatomy and neural networks. Here the observed changes in glomerular volumes, density, innervation and neuroconnectivity in cortico-medullary axis of key FTUs, across ages and conditions uncovers how mature function and homeostasis may be maintained in the kidney (see below).

One novel intriguing discovery from our work is the observation that nerves connect glomeruli in a community and across communities through secondary connections encompassing the glomeruli and between glomeruli by both sympathetic and sensory nerves. These further connect to other nephron segments including various tubular FTUs indicating that there may be unrealized complex feedback mechanisms among nephrons beyond the sensory-sympathetic regulation at the vascular pole ^19^. The secondary connections caused by nerves meandering between glomeruli may invoke a coordinated response in a connected community by amplifying or dampening activity compared to a “lone” glomerulus.

Our discovery and introduction of the concept of mother glomerulus connecting glomerular communities suggest that they may serve as important local control centers, similar to the hubs of spoke-and-hub networks ^26^. For example, an increased density of sympathetic fibers may make a mother glom more fine-tuned in responding to and modulating changes in glomerular blood flow and blood-pressure. Serving as an intersection point for sensory fibers from multiple communities may allow mother glom to serve as a highly sensitive and efficient control centers than if every glom was equally involved in the network ^26^. Our data supports a scale-free network as mentioned above indicating that this design in kidney makes it less prone to random failure of nodes in the network where the mother glomeruli serve as key hub points. Being a scale-free network would importantly make the neuronephro-network resistant to the accidental failure ^27^ of FTUs due to accumulating sclerosis through age or minor damage, as the loss of less neurally-involved gloms would prove largely inconsequential to network behavior.

These observations lead us to provide interpretations for several clinicopathological observations. For example, the effectiveness of denervation relieving symptoms in diseased kidneys ^5^ indicates these networks may frequently reach a point of maladaptive failure in age and disease. Our images demonstrate perturbation of the neural network in both age and disease, through incorporating sclerotic components within unusually well-connected networks, or under-connected networks where nerves have retreated from diseased regions. A case may be made that the abundance of spokes allows the networks to sufficiently compensate for glomerular loss until random processes have led to the disincorporation, or instead decoherence through over incorporation, of mother glom hub points. When this threshold is crossed, the network would cease to function as intended, with further neural signaling confounded by dysfunctional network components to the point of maladaptation, worsening disease states and creating a vicious cycle that further damages the network. By completely removing the network through denervation, the kidney is reduced to a uninnvervated, auto- regulated state, potentially improving symptoms through the elimination of a bad network. The persistence of symptomatic relief after reinnervation ^5^ may consequently be attributed to the new network “resetting” itself from bad components, reassembling itself around remaining FTUs in a structure of proper functioning.

Our observation of diminishing neuroglomerular connectivity in aged, disease or neonatal samples may provide an explanation for changes in eGFR and relation to hypertension due to variations in nephron number. With evidence of fewer glomeruli at birth being associated with diseases like hypertension ^37,38^, this does raise a question of whether a similarly diffuse neural network is a factor for long-term kidney outcomes. It may also be the case that low density glomeruli simply do not require as extensive of a network to regulate due to their gross reduction of parallelized components, where diffuse networks would be hypothetically appropriate as glomeruli become less dense with age ^35^.

We recognize several limitations of our study. 3D imaging methods for solid organs are still being improved and clearing methods in conjunction with LSFM are limited in multiplexity, have long experimental time exceeding 3 weeks and show variability in antibody penetration and staining. Further, uniform deeper tissue imaging is challenging with increasing depth in LS microscopes. While we present several observations based on thousands of nephrons, often from two or more samples per patient, in future expanding this to more diverse group of patients is needed. Nevertheless, we were able to discover important trends across age trajectory for several of the observations several consistent with previous works.

Overall, the 3D maps of the human kidney in micro and macroscales of several key FTUs and their relationship to neurovasculature and organization into neighborhoods and communities and transitions at key postnatal time point in health and disease provide previously unappreciated insights into how structure informs on function and its impact on homeostasis, disease and therapy.

## METHODS

### Statistics and Reproducibility

For 3D LSFM imaging experiments, antibodies selected were commercially available, validated by confocal microscopy, several FOVs were imaged from each sample and in several cases more than 1 sample was used for same patient **(Extended data Supplemental Table 1).** For adult kidneys, 2 or more patients were in each age group with >5 FTUs analyzed.

The following statistical tests were used to establish significance:

1. Two sample Welch’s t-test for all comparisons with n > 5 was used to establish significance between cohorts. Welch’s t-test is an alternative to Student’s t-test that offers greater power when population variances cannot assumed to be equal ^39^.
2. Two sample Wilcoxon rank-sums test was used when n = 5 (only the confocal datasets) ^40^, due to offering greater power when t-test assumptions of normality and variance cannot be met due to low n.
3. Kolmoglorov-Smirnov test for normality was used to assess possible non-normality of distributions n ≥ 50. ^41^

### Ethical Compliance

We have complied with all ethical regulations related to this study. All experiments on human samples followed all relevant guidelines and regulations. For pediatric samples, human tissue was obtained through the non-profit United Network for Organ Sharing facilitated by the International Institute for the Advancement of Medicine and processed into the University of Rochester LungMAP .BRINDL biorepository before transfer to the Kidney Translational Research Center (KTRC) for use in the Pediatric Center for Excellence in Nephrology at Washington University. Next of kin authorization for research was obtained for all donations. The project is overseen by the University of Rochester Research Subjects Review Board as non- human subjects research protocol authorized for the use of the de-identified tissue from the deceased donor (RSRB00056775). Adult reference kidney samples as part of the Human Biomolecular Atlas Program (HuBMAP) consortium and disease kidneys were collected by the Kidney Translational Research Center (KTRC) under a protocol approved by the Washington University Institutional Review Board (IRB #201102312). Informed consent was obtained for the use of data and samples for all participants at Washington University, including living patients undergoing partial or total nephrectomy or from discarded deceased kidney donors.

### Sample Procurement and Preservation

The process for procuring pediatric tissue has been reported [dx.doi.org/10.17504/protocols.io.bjuxknxn]. Briefly, kidney was handled as transplant quality, stabilized in transplant buffer solution, and referred for research as unable to place for transplantation. Adult kidney samples were obtained from deceased or surgical nephrectomy donors as reported {dx.doi.org/10.17504/protocols.io.8epv5r64jg1b/v1; dx.doi.org/10.17504/protocols.io.261ge56zdg47/v1}. An ∼2cmx3-5mmx1-3mm slice of kidney was dissected from a renal lobe, typically containing both cortex and medulla and fixed with 4% paraformaldehyde (PFA) overnight at 4 °C and stored in phosphate buffered saline (PBS) with preservative until use.

### Tissue Clearing and Blocking

A detailed, step by step procedure for clearing through imaging is available at dx.doi.org/10.17504/protocols.io.eq2lyjxmqlx9/v1 After fixation, samples were cleared using the CLARITY SHIELD Active clearing protocol ^12^. First, samples were immersed in 20 mL of SHIELD-OFF solution (10 mL LifeCanvas SHIELD-epoxy, 5 mL LifeCanvas SHIELD-buffer, 5 mL de-I water) for 3 days at 4°C. Next, samples were immersed in 20 mL of LifeCanvas SHIELD-ON reagent with shaking for one day. Finally, samples were immersed in LifeCanvas delipidation buffer and underwent active clearing using the LifeCanvas Smartbatch+ delipidation chamber, enhancing delipdation effectiveness and speed. The samples were washed twice with in 1x PBS with rocking for three hours each, then once more overnight. Washed samples were blocked with 1%BSA/0.2%skim milk/0.3% Triton X-100 in 1x PBS (PBS-BB)/0.1% Sodium Azide at room temperature (RT) for 2-3 days with rotation.

### Immunofluorescence Staining

Details of the antibodies used are provided in Supplementary table 1 & 2. Glomeruli were stained with the anti-NPHS1, an antibody marker for nephrin that labels podocytes in the glomerular tuft ^42^, or anti-PODXL which labels podocytes and glomerular capillaries among other vessels. Collecting Ducts were stained with anti-AQP2, labels principal cells in the collecting duct. Nerves were stained with anti-Tuj1, an antibody that labels TUBB3. Endothelial cells of the blood vessels were stained with the anti-CD31 or anti-CD34 antibodies. Sympathetic nerves were stained with the anti-TH labeling tyrosine hydroxylase. Sensory nerves were stained with anti-CGRP. Synaptic termini for the nerves were stained with anti-Synapsin 1. Thick ascending limb was stained with anti-UMOD. Proximal tubule was stained with anti-LRP2. Primary antibodies were used at 1:100 or 1:200 dilution. Secondary antibodies used were anti-mouse alexa-488, anti-rabbit-cy3 and anti-goat-cy5 and anti-sheep-cy5 and at 1:150 dilution. Nuclei were stained with DAPI at 1:1000 dilution.

Samples were incubated with primary antibodies in PBS-BB with rotation either at room temperature for seven days at room temperature or at 37°C for 2 days, then at room temperature overnight. Samples were removed from primary and washed with 0.3% Triton X-100 in 1x PBS/0.1% Sodium Azide at room temperature with rotation overnight with several changes followed by incubation with fluorescence-conjugated secondary antibodies in 0.3% Triton X-100 in 1x PBS/0.1% Sodium Azide at room temperature with rotation for 2-4 days. Next, samples were washed with 0.3% Triton X-100 in 1X PBS/0.1% Sodium Azide at room temperature with rotation with several changes overnight at RT. Samples were incubated with DAPI 1:1000 in 0.3% Triton X-100 in 1X PBS/0.1% Sodium Azide at room temperature for one day to label nuclei. Finally, samples were washed with 0.3% Triton X-100 in 1x PBS/0.1% Sodium Azide at room temperature overnight at RT; DAPI could also be included during incubation with the secondary antibody. After staining, the sample were stored in the dark, wrapped in aluminum foil in 1x PBS/0.1% Sodium Azide prior to imaging. We typically previewed the sample using confocal microscopy before LSFM. When ready for LSFM, the sample was refractive index matched with LifeCanvas EasyIndex. The sample was transferred into a new 5mL Eppendorf tube and incubated sample at RT in a 50/50 mixture of EasyIndex solution for one day. Next, the sample was incubated in EasyIndex at RT for one day.

### Imaging using LSFM

All samples were imaged on the ZEISS Lightsheet 7. A small amount of super glue was used to affix the sample to a glass coverslip, which was mounted in an imaging chamber filled with 1.52 refractive-index Cargille Oil. First, the entirety of the sample was imaged with a 5x objective with a zoom ranging between 0.4 to 1.0x to capture the entire sample, depending on sample size. The 5x image was used to identify several regions of interest that were imaged with a 20x resolution lens and dual 10x illuminators at 1.0 zoom, using a 2x2 tile grid and imaging 1mm deep in Z-axis, resulting in 20x field of views (FOVs) around 1mm^3^ on average. All images thus specified were taken using Cy5, Cy3, 488, and 405 laser lines. Additional large 20x FOVs up to 8mm^3^ were imaged on select samples to evaluate the relationships of nerves and nephrons at mesoscale at high resolutions; here data was instead imaged on two or three of the four laser lines to facilitate data processing and make these large FOVs conducive for analysis.

### Data Processing

#### General Data Processing Pipeline

The general pipeline to transform raw image data from the Lightsheet 7 into 3D isosurfaces involved the following steps.

##### 1. Stitching

The Lightsheet 7 microscope captures large volumes by creating overlapping Z-stacks, requiring stitching into a singular image. For 5x volumes and 20x volumes below 500 GB, the stitching was with the stitching function in ZEN Blue 3.3. For 5x volumes and 20x volumes above 500 GB, the stitching software Stitchy was used to stitch the volumes.

##### 2. Down sampling

To enable efficient data processing and analysis, both 5x and 20x volumes were down-sampled 4x in both the X and Y dimensions, resulting in a 16x down sampling. For images stitched in ZEN Blue, the ZEN downsampling algorithm was applied on stitched files by setting pixel sizes in X and Y to 0.25. For images stitched with Stitchy, down sampling occurred concurrently, generating a secondary file with X and Y pixel sizes expanded 4 times.

##### 3. Masking

Segmentation through the creation of binary masks reduces an image to solely positive or negative signal, enabling efficient computation. For this study, binary masks were generated utilizing either Cellpose2 or Cellpose 3 ^43,44^ or the ImageJ extension Labkit ^45^. Cellpose is a trainable machine learning software that allows for custom model segmentation. Models were constructed by manually segmenting 2D slices from various LSFM datasets. A model to segment anti- NPHS1 labelled Glomeruli was constructed from 179 images across 6 20x datasets, and 16 images across 4 5x datasets. A model to segment anti-AQP2 labelled collecting ducts was constructed from 46 images across 3 20x datasets. The model to segment DAPI labelled nuclei was constructed from 3 images across 2 datasets.

These models were trained from scratch based on the dataset with 10,000 training epochs each.

Labkit, an extension for ImageJ, allowed trainable machine learning segmentation of heterogenous objects in lightsheet volumes. A custom model was trained from scratch for each FOV segmented. Labkit was trained to segment channels containing filamental objects including Anti-Tuj1 labelled nerves for 5x and 20x volumes, anti-CD31 or anti-CD34 labelled vessels for 5x and 20x, and labelled tubule structures such as CDs, PTs, and TALs for 5x.

Files created by both cellpose and labkit were saved as grayscale TIFF files. These masked TIFFs were converted to 8bit binary in FIJI or Python, setting all nonzero signal to the maximum 8bit value of 255.

### 4. Thresholding and Isosurface Generation

To convert masked volumes into isosurfaces in Imaris for both visualization and analysis, binary TIFF masks were first converted to .ims files using the Imaris File Convertor Program, inputting the appropriate X, Y, and Z voxel dimensions. Binary channels were then imported into Imaris overtop the stitched, down-sampled volumes. Since binary channels separate objects into signal or background, all binary channels had the least possible intensity threshold of 0.125 applied onto them, removing no pixels from the mask to isosurface.

Surface smoothing was also disabled for all isosurface generation. However, voxel-size based thresholding of objects was applied as needed to remove any small, mislabeled masks from the isosurface.

### Qualitative/Quantitative Analysis of Volumes

#### Segmenting Structures in 5x for Visualization and Analysis Segmenting Glomeruli

Glomeruli in 5x were segmented with their 5x Cellpose 2 model and turned into an isosurface in Imaris.

An additional step during isosurface generation involved using the Imaris surfaces seed separation method, which allowed the separation of adjacent fused objects by morphological size. Due to the thickness of the light sheet, often empty space regions just adjacent to the tissue yielded a low intensity signal where there should have been none. These “glomeruli shadows” would sometimes be selected during segmentation as positive signal. To obtain accurate glom counts and volume information, these incorrectly selected objects were manually removed from the resulting surface within the Imaris isosurface. The resulting glomeruli isosurface was used to mask the original binary glom channel. This newly masked channel was exported for analysis.

#### Segmenting CDs, TALs, and PTs

Tubule structures in 5x were segmented using a Labkit model and converted into an isosurface in Imaris. Since we took advantage of region specificity of glomeruli and CD and stained them with the same secondary antibody, for CDs, NPHS1 glom signal was removed using the 5X glomeruli isosurface to mask the tubule segmentation if necessary. The resulting masked tubule channel was exported as a tiff image for analysis.

#### Segmenting Nerves

Nerves were segmented using a Labkit model, then traced in Imaris using the Filaments tool, which automatically converted the segmented nerve mask into a more cohesive filament network. The resulting filament network was converted into a binary channel using the Imaris Convert Filaments to Channel Xtension and exported as a tiff. This represented a refined nerve mask for 5x that was used for network analysis.

#### Segmenting Vessels

Vessels in 5x were segmented using a Labkit model. Due to the flexibility of labkit segmentation, vessels in 5x were either segmented for CD31/CD34 signal, which returned a mask displaying the vessel wall stain, or they were segmented for empty space within the vessels, which returned a mask displaying the entire volumes of larger vessels. The latter approach was used to both better visualize vessels as 3D objects, and to account for inconsistent CD31/CD34 staining. Masks segmented for empty space within vessels were converted into an Imaris isosurface and had any erroneously selected regions, often at the edge of the tissue, manually deleted. Vessel masks in 5x were only used for visualization and not quantitative purposes.

### Quantitative analysis in 5x images

#### Basic Sample Measurements – Volume, Length, Width

Basic sample measurements were obtained using Imaris. The volume of an entire sample was obtained using the Imaris surfaces tool. First, a temporary, down-sampled image was created to increase processing speed. Next, a new surface was autogenerated on the 488 channel of the 5x image. With smoothing disabled, this surface was thresholded until the surface enveloped the entirety of the tissue volume but none of the background. The volume of the resulting surface was obtained from Imaris’s surface statistics tool **(Supplemental Table S11)**. The surface was also used to generate a binary mask to represent the spatial properties of the entire sample.

The length and width of the sample was obtained using the Imaris measurement points tool. The cortical length and medullary length of the sample were also obtained with the measurement points tool, measuring from the top of the cortex to the corticomedullary junction, and from the corticomedullary junction to the base of the medulla.

#### Determining Glomerular Count and Density in the Cortex

Glomeruli counts of an entire sample was estimated by 3D watershedding the 5x analysis glomeruli masks. Watershed ^46^ is an image processing method for separating fused binary objects. We used the watershedding algorithm “Distance Transform Watershed 3D” from the FIJI plugin MorphoLibJ ^47^ a function specialized for separating binary objects in 3D. This method estimated glom count with 97% accuracy in sample SK5 compared to manual count, for example.

Glomeruli density in the cortex specifically was estimated using a script written in Python. This was accomplished by dividing the total number of glomeruli by the volume of their convex hull. Since glomeruli appeared in cortical regions but not medullary ones, a volumetric boundary that surrounded only regions with gloms was necessary to calculate density. A 3D convex hull ^48^ was generated around the analysis glom mask using the scipy.spatial module ConvexHull. This represented the smallest convex 3D polyhedron that enveloped all binary signal, which was used to represent the volume of all glomeruli-occupied space. Dividing the number of glomeruli by the volume of the convex hull thus yielded the number of glomeruli per unit of 3D space in the cortex. **(Supplemental Table S11).**

### Quantifying Glom-Nerve Interactions

The neural innervation of glomeruli was procedurally quantified by skeletonizing ^49^ the Tuj1 nerve mask in FIJI, followed by using a Python script to calculate an average, normalized density of nerve skeletons within 10 μm of glomeruli. The Tuj1 binary mask was used over the filaments nerve mask since it represented a more accurate description of the size and shape of the nerve fibers, and responded better to skeletonization, while the nerve mask was better utilized for establishing connectivity specifically. First, Tuj1 masks were skeletonized in FIJI using the plugin Skeletonize (2D/3D). This tiff file was converted to a 3D numpy array alongside the analysis glom masks.

To enable accurate comparative analysis across images taken at differing resolutions, all arrays underwent a resolution normalization step where the scipy zoom() function re-interpolated the glomerulus and nerve-skeleton arrays to all have the same voxel sizes as one another. In essence, low-resolution arrays were “stretched” until their voxel sizes were the same as the smallest reference array in the dataset. The interpolated nerve skeletons were then skeletonized once more with the skimage skletonize_3d function. This was especially important for 5x images taken at separate zooms, as normalization allowed all nerve skeletons to represent the same magnitude of “nerve” across all volumes being compared.

The glomerulus masks were then dilated by 10 µm in all dimensions using the OpenCV dilate function. Next, the non-dilated glom array had its indices inverted, turning 255 to 0 and 0 to 255, which was multiplied against the dilated glomerulus array to return an array of all regions within 10 µm of a glomerulus. The total pixel count of this search-region array was likewise obtained. In 5x **(Supplemental Table S6/11),** the dilated binary glomerulus array was multiplied against the nerve skeleton array to mask out all nerves not within 10 µm of the glomerulus. In 20x **(Supplemental Table S7)**, rather than using dilated glomerulus to mask nerves like 5x, instead the search region itself was multiplied against the nerve skeleton array to mask out both all nerves not within 10 µm of gloms and all signal within the tuft. This differing strategy between 5x and 20x was utilized because the majority of any nerve signal within gloms tufts was noise at 20x, but not at 5x, due to the resolution differences. The number of remaining white pixels in the masked nerve array was then calculated.

The density of nerve skeletons around a glomerulus in the volume was finally calculated by dividing the pixel count obtained for the nerves by the pixel count obtained by the search region. Ultimately, this density was the value used for comparative analysis. This script also returned an estimated true search volume and total glom search volume by multiplying the search volume and glomerulus masks by the normalized 3D voxel sizes. For simple comparison between broad sample categories (such as reference vs diseased), the automated neural data was also aggregated across FOVs by calculating for each an average weighted by the total number of glomeruli analyzed in a sample. First, the total number of glomeruli analyzed in a sample was estimated by dividing the total volume of gloms analyzed in that sample by the average glomeruli volume of that sample calculated from 20x volumes **(Supplemental Table S12)**. This value was used to obtain an aggregate mean of a category by first finding the total number of estimated glomeruli in that category (such as total number of all reference gloms across all samples), then for each sample in said category, dividing that sample’s estimated glomeruli analysis count by the total estimated glom analysis count and multiplying this ratio by that sample’s average glomular nerve density score. Adding up every value for each sample in a category obtained this way produced a single aggregate estimate for the glom-neural density of that category.

The neural innervation of glomeruli was manually quantified by tracing the filaments within 10 µm of gloms using Imaris **(Supplemental Table S8).** First, only gloms fully contained within an FOV were selected and used to create a binary mask, which was exported as a tiff. These gloms were dilated by 10 µm in all dimensions using Python (OpenCV) to establish a boundary region. This dilated tiff was imported back into Imaris. All nerves in the dilated regions were manually traced from the Tuj1 channel in Imaris using the Filaments function. The lengths of nerves around a particular glom were then determined by selecting only that glom’s nearby traced filaments in Imaris and finding the summated segment length in the Imaris statistics tool. The volume of dilated gloms in the FOV was calculated by converting them into an isosurface in Imaris and exporting their volumes through the statistics tool. Innervation density in an FOV was then determined with the following formula: (Total dilated glomerulus vol – glomerulus vol)/Total nerve length of glomeulus.

### Determining Densities of Structures Across Regions of Tissues

The density of structures across a tissue sample were obtained by using Python. This was accomplished by slicing through the masked 5x data in the direction of the Y axis and dividing the total voxel count returned by np.count_nonzero() by the voxel count of the binary mask. This outputs for each plane were saved in excel as two columns representing depth in the tissue and the density of that plane, respectively.

Glomeruli and CD density distributions sometimes contained extreme density outliers obtained from regions towards the top and the bottom of the tissue totally occupied by a glomerulus **(Supplemental Table S4)** or duct **(Supplemental Table S5).** These outliers were removed using the equation Outlier = Q1-(1.5(IQR)); Q3+(1.5(IQR)), with Q1 representing the data’s first quartile, Q3 its third quartile, and IQR the value of Q3 - Q1. Nerve densities were grouped into buckets of n=5. In most instances, samples were imaged from cortex to medulla in the y-axis. Consequently, these distributions were used to analyze how densities changed through cortical and medulla depths. However, differently oriented images were not used for this specific analysis purpose. Planar distributions for nerve densities **(Supplemental Table S6)** were used as a surrogate measure general innervation distribution across a sample, as shown in the innervation violin plots of Figure 5.

### Coregistration of 20x onto 5x to Measure Cortical Depth

20x FOVs were coregistered onto 5x volumes using Imaris. Due to the Lightsheet 7 requiring switching immersion chambers between 5x and 20x imaging sessions, all 20x FOV regions had to be coregistered post-imaging, rather than in-tandem. While imaging 20x volumes, the gross location of the 20x image was first labelled on a 2D image of the 5x volume to act as a rough guide for realigning the 20x. After stitching and downsampling, the 20x FOVs had their 647 channels extracted, converted into an ims file, and imported into Imaris atop the 5x image. Using the labelled 2D 5x image as a reference, Imaris was used to rotate and translate the 20x FOVs into their correct locations within the 5x ims file, using glomeruli as fiducial markers.

Since this coregistration was accomplished manually, it did not result in exact pixel-to-pixel correspondence, but did approximately describe values such as cortical depth of a 20x FOV. After each coregistration, the cortical depth of a 20x FOV was obtained using Imaris’s measurements points tool to measure the depth from the top of the cortex of the tissue slice in 5x to the center of the 20x FOV. **(Supplemental Table S16).**

### Segmenting Structures in 20x for Visualization and Analysis

#### a. Segmenting Glomeruli

20x glomeruli were roughly segmented in Imaris with the spots tool, then finely segmented in Cellpose, then manually refined using the Imaris surfaces tool. The stitched, downsampled 20x volumes were first converted into Imaris images. Within Imaris, regions containing glomeruli were first manually labelled using the Imaris spots feature, which allowed the masking out of larger spherical regions containing gloms, which represented a rough segmentation to aid in computationally locating gloms. The use of spots proved to increase segmentation accuracy by acting as a queue to the model that a glom would be inside a spot, however this was skipped for certain high-density images that had sufficiently strong signal. The masked glomerulus file was exported from Imaris as a tiff and segmented using its Cellpose model, with the mask then converted into an isosurface within Imaris. In a few instances where NPHS1 stain was weak, nearby areas of tubules or ducts would occasionally be masked adjacent to a glom mask, causing the two to fuse in the isosurface. These incorrectly segmented regions were manually cut off and deleted using the Imaris surface cut tool. Within the isosurface, any nearby glomeruli that erroneously fused were also cut apart manually along one clipping plane with the surface cut tool. This represented the analysis glom isosurface used to quantify glom morphologies.

To accurately characterize glom-nerve networks, a second segmented glomerulus mask was created that represented both an accurate segmentation of gloms, but also had no touching regions between adjacent gloms. To accomplish this, two clipping planes were used to split the junctions of any fused gloms within the analysis isosurface with the Imaris surface cut tool, and the resulting, small in-between region was deleted.

This edited isosurface was used to mask over the original 20x binary gloms. The resulting masked glomeruli were exported for analysis. In certain high-density glom volumes, this was instead accomplished algorithmically by first downsampling the 20x volume, a step that all glom files underwent anyway while creating 20x networks, followed by watershedding using MorphoLibJ’s “Distance Transform Watershed 3D” ^47^.

#### b. Segmenting Tubule Structures

CDs were segmented by their Cellpose model and converted into an isosurface in Imaris. PTs and TALs, which did not have a Cellpose model trained, were instead manually segmented within Imaris using the filament tracer tool to trace the inside of the tubule, for visualization and qualitative analysis when relevant.

#### c. Segmenting Nerves

Nerves in 20x volumes were segmented using a Labkit model then traced in Imaris using the Filaments tool, which automatically converted the segmented nerve mask into a more cohesive filament network. The resulting filament network was converted into a binary channel using the Imaris Convert Filaments to Channel Xtension and exported as a tiff. This represented a refined nerve mask for 20x that was used for network analysis.

#### d. Segmenting Vessels

Vessels in 20x volumes were segmented using a Labkit model. These could be converted into either a filament network or isosurface in Imaris, however the vessel segmentation was used only for visualization and not quantification, due to its inconsistent stain.

#### e. Segmenting Nuclei

Nuclei in 20x volumes were segmented using their Cellpose model. Only binary glomeruli fully contained within the FOV were used to mask the 405 channel in Imaris. The resulting masked channel was exported as a tiff, which was used to segment for nuclei. As the high frequency 405 laser did not penetrate deeper tissue as well as the other laser lines, oftentimes only gloms at the surface of an FOV were included in this analysis, while those deeper in the tissue were excluded due to their reduced signal.

### Quantifications In 20x

#### Glomerulus Morphology

Glomerulus volume and sphericity was obtained from the glomerulus isosurface in Imaris. Only glomeruli contained entirely within the 20x volume were selected within the glom analysis isosurface. The volume and sphericity for those gloms was exported from the Imaris statistics tool. **(Supplemental Table S12).**

#### Glomerulus Cell Counts

Cell counts in a glomerulus were estimated using Cellpose 2 segmentation. The segmented mask obtained from cellpose maintained identities of individual cells by labelling them with separate grayscale values. The number of cells within an individual glom could therefore be obtained by first using Python to label the analysis glom masks with separate grayscale values as well. By iterating through all labelled gloms, each could be converted into a boolean mask that when multiplied against the cell array would return an array only containing all cells within one glom. These cells were then counted by flattening their 3D array, removing any zero values, and converting the resulting list into a unique set whose length was equivalent to the number of unique objects. Cell densities could likewise be obtained by dividing this value by the volume of its corresponding labelled glomerulus object. **(Supplemental Table S15).**

#### Determining the Orientations of Glomeruli in 3D Space

Glomerulus orientation was categorized by the angular direction in which its vascular pole was oriented in 3D space using Imaris to label the location of vascular poles, followed by using a Python script to measure angles. First, the gloms separated for network analysis had the locations of their vascular poles labelled in 3D space using Imaris’s spots feature. A python script was used to generate a completely white volume of the same dimensions as the channels in Imaris. Next, Imaris’s surface mask feature was used to create binary masks of only the analysis glom isosurfaces fully contained in the FOV. The spots at the vascular poles were also used to mask over the white volume to create a tiff containing only binary spots denoting the locations of vascular poles. Both of these binary channels were exported as tiff files and downsampled in FIJI to speed up processing.

Next a Python script was used to obtain angular information about the glomeruli from the exported channels. The script first calculated the centroids of all vascular pole spot markers from the pole tiff and matched them to the nearest glomerulus object in the separated glom mask. The centroid of this glomerulus was calculated and both pairs of centroids were matched. The line between these centroids was treated as a surrogate for the 3D orientation of these gloms, with the vascular pole spot centroid as the end point and the glom centroid as the base point of a vector. First, the azimuth angle was calculated with the numpy method np.arctan2(dy/dx), while the polar angle was calculated with np.arctan2(dy/dz). Note that np.arctan2() function is programmed to specially accommodate arctan calculation in all four angular quadrants. The resulting angles for each glom/pole pair were exported in an excel file. **(Supplemental Table S3).**

To visualize the orientations of gloms, verify correct centroid matching, and search for common fates, the centroid pairs were further used to draw rays from the glom centroid, through the pole centroid, to the end of the volume. This was done using a python script that first used the slope of the vector to find the furthest allowable point in 3D space away from the glom centroid, before using the command np.linspace() to estimate a set of linear coordinates of the vector within the array and set these indices to white. A tiff file was exported with all the drawn rays, which could be opened over the original channels in Imaris to easily view exactly how gloms are oriented in 3D. **(Supplemental Figure 1Sh).**

#### Validating Automated Neural Innervation Analysis with Manual Analysis

Manual and automated analysis demonstrated similar trends between FOVs. To examine this, same FOVs were used for both analyses. For example, manual and automated analysis of the same FOVs throughout samples SK1 and SK2 [PPID: 3785] yielded a shallow linear trendline, with the automated trendline a little less positive than the manual one. For further validation, this comparison was normalized by adding the difference between the manual and automated SK1 FOV2 calculations to all the automated measurements.

This was necessary to bring automated measurements into a generally similar magnitude range as manual measurements, since automated measurements relied on a different calculation. A two-sample Welch’s t test between these manual and normalized automatic distributions failed to reject the hypothesis that these distributions are different (p = 0.761). While not entirely perfect, the automated pipeline allowed efficient comparison of innervation magnitudes that was representative of true biological structure, trading a small amount of accuracy for a massively increased sample size. **(Supplemental Figure 1Sd).**

#### Generating Neuroglomerular Networks

To enable this analysis, we developed “NetTracer3D”, a novel Python tool able to generate large, undirected networks from 3D images, in addition to providing analytical support. With highly customizable parameters, NetTracer3D offers several derivatives of its core algorithm to allow network generation for a variety of circumstances, accuracies, and speeds. Two versions of NetTracer3D were developed, utilizing the same core logic, with V2 offering superior customizability and computational performance on larger datasets. The majority of analysis in this study was accomplished using V1, except for the large 20x dataset shown in Figure 4d and the AKI 5x dataset in Supplemental Figure S3b.

### The Core Algorithm

Both versions of NetTracer3D used the same core logic. Glomerulus and nerve masks were designated as nodes and edges in a network, respectively. In general, nerve masks existed as one or a few interconnected objects in the volume that if utilized directly to establish a network would always result in erroneous connections between the majority of nodes. To establish discrete node-node connections, the node masks were first used to discretize the network. This was accomplished by allowing the nodes to extend outward in a 3D-search space and break any edges passing through their space into separately recognized connections. Essentially, a nerve passing over a glomerulus would be split by the glom’s search space into a before and after segment. By differentiating edges in this manner, distinct connections between objects could be recognized within the neural network. Connections could also be made through edges retained within the search spaces of two neighboring nodes. Any connected objects were then saved in the network as a node pair, which could be analyzed using the Python NetworkX package ^50^. As this algorithm is designed to function for any generic 3D image containing potential node and edge objects, it additionally provides future analytical use beyond this study.

### NetTracer3D V1 Specific Implementation in This Study

#### Establishing Nerves as Edges (V1 & V2)

The processed glom and nerve binary masks were converted to 3D binary numpy arrays in Python.

Since the nerves wrap the outer boundary of the Bowman’s capsule rather than entering inside the glomerulus, first the glom array was dilated by 10 µm in all directions to allow each glom to envelop its innervating nerves. After enveloping nearby nerves, these gloms were used to mask out that region, effectively splitting up the nerve network into a series of glom-to-glom neural connections, rather than one single interconnected bundle. This was accomplished by inverting the indices of the dilated glom arrays, marking the dilated gloms as black null regions and everything else as white regions. The inverted array was multiplied against the nerve array, removing any nerve segmentations that came within ∼10 µm of gloms. This allowed the nerve network to be broken up into subconnections between gloms, whereby the separated nerves served as edges in a network, and the gloms as the nodes. The resulting array contained the “outer edges”. Due to the fact that nearby gloms sometimes had nerve connections between one another that were completely enveloped by the dilated regions and therefore eliminated from the edge network, a second set of edges was created by multiplying the non-inverted dilated glom array by the nerve array. The resulting array contained the “inner edges”.

#### Optionally Removing the Nerve Trunk (V1 & V2)

To examine subcommunities of gloms in absence of the nerve trunk that provided extensive network innervation, at this point the nerve trunk could be removed. This was accomplished by identifying the largest object in the outer edges network and setting its indices to black. For consistency, all volumes analyzed for their glom-nerve networks were processed twice, both with and without the trunk, to compare the analytical results.

#### Downsampling, Labelling, then Dilating Edges (V1 20x Only)

Due to being an earlier implementation, NetTracer3D V1 utilized a greedier algorithm that required the user to perform an intermediate downsampling step at 20x resolution. The purpose of this step was to prepare the glomerulus array for an ID-specific dilation into each other’s search spaces, but also served as necessary to allow nerves whose segmentation was not a complete filament to become reconnected. In V2, this reconnection is accomplished via a dilation algorithm within the edge array, but in V1, downsampling accomplished an equivalent goal, essentially “squishing” punctated nerve masks back together.

Using V1, both sets of edges were saved as 8-bit tiff files. Each edge tiff was opened in FIJI and had a maximally brightening look up table applied. The resulting image was downsampled in FIJI by 5 in all dimensions using the scale function with default settings. The downsampled edge images had a maximally brightening LUT applied before they were saved. The reason for downsampling in FIJI instead of Python is that this specific set of steps in FIJI will downsample the binary network, yet prioritize retaining positive indices that represent nerve signal. By comparison, the scipy downsample method will maintain the ratio of positive to negative elements while downsampling by eliminating positive indices, losing network coherence. This FIJI downsampling method effectively allowed regions of the network that had broken up into punctates due to imaging or segmentation artifacts to reconnect to one another without the need of a large dilation/closing step. Both sets of downsampled edges were then opened in python as 3D binary numpy arrays. Both edge files were labelled, a scipy method that assigns unique numerical identities to all separated objects within the array. Finally, the labelled array containing the outer edges was dilated by a single pixel in all dimensions, which helped further reconnect any broken regions of the network while also allowing it to overlap the dilated gloms that had previously been used to mask over the nerve network. This overlap was crucial in identifying which edge interacted with which gloms, while the single pixel dilation in the downsampled file proved relevant in reconstituting network loss due to imaging artifacts. Several downsampling and dilation parameters were tested, with the above steps best demonstrating a strong ability to accurately identify connections based on comparative observations with the 20x images themselves.

#### Downsampling and Dilating Labelled Gloms into a 3D List Array (V1 20x Only)

The glomerular array was labelled to give each glom a unique numerical identifier. The glom array was downsampled in all dimensions by 5, resulting in an array with the same dimensions as the downsampled edges. Next, a new array of the same dimensions as the downsampled glom array was created, containing an empty list in every index. An iterative dilation method was used to dilate each labelled glom by 10 µm in every dimension within the list array. Using an array with list indices allowed dilated glom regions that overlapped to preserve their respective identities within an index, rather than a standard dilation method which would instead force one dilated glom to override another in the instance of overlap. Note that this specific method was later replaced in V1 with an improved subarray method designed for V2, which accomplished the same exact computation at increased speeds.

#### Establishing Pairwise Connections in the Network (V1 20x Only)

After the above downsampling, the labelled edges and labelled gloms were ready to have their network connections established. For the 20x networks, this was simply accomplished by iterating through the edge indices and compare them against the labelled glom array to see which discrete nerves interacted with which gloms. Edges that overlapped with two or more glom labels were used to generate a list that contained the numerical identifiers of all gloms connected by the edge. This list was added to a master list categorizing all connections in the network. This process was repeated on all edges in the network. The concurrent.futures Python module was used to parallelize this step and to further increase processing time. Overall, this method was computationally intensive but more than sufficient to process the downsampled 20x arrays.

#### Establishing Pairwise Connections at 5x with High-Speed Array Trimming (V1 5x and V2)

Due to the lower resolution of 5x volumes, downsampling and dilating the nerve volume was not necessary to reestablish missing connections, and would only cause the loss of network complexity. Similarly, dilation of gloms into overlapping lists was far too slow for the number of gloms in the volume, but lower resolution meant overlapping detection neighborhoods was less of an issue. As a result, the V1 script was modified to be faster and more streamlined for 5x analysis. Samples were first dilated by 10 µm to establish detection neighborhoods using OpenCV. Next, overlapping gloms were separated using MorphoLibJ’s “Distance Transform Watershed 3D” ^47^. This method preserved the correct identities of dilated, separated gloms in 5x with 95.4% accuracy versus manual separation in sample SK5. Note that this dilation followed by re-watershedding was no longer necesary in V2 due to the implementation of smart-dilation (see NetTracer3D V2 improvements). In V1, the dilated, watershedded glom masks were separated and labelled with separate grayscale values index by the plugin, and therefore could be used directly to create networks. First, the dilated glom mask was used to break the network into discrete units in the same manner as for 20x volumes.

However, due to already being dialted into a search region, the 5x script did not require listwise dilation, glom labelling, or downsampling. The separated nerve network was still dilated by a single pixel in all dimensions, both to address any missed connections and to allow overlap with glom search regions.

For identifying pairwise connections at 5x, a computationally efficient method was developed to allow fast processing in large arrays with hundreds, or even thousands, of gloms. This was accomplished by using array transforms to create a 3D mask for both the glom and edge arrays that only contained labelled components in each that overlapped with one another. Using these masks, any components of the arrays that were not overlapping could be set to 0 with one computational step. Next, these arrays were flattened into 1- dimensional lists and had any indices containing a 0 removed, significantly trimming the number of indices to search through. Finally, the shortened edge list could be compared to the glom list. All values in the edge list of the same numerical label represented the same nerve spatially, which when compared to the glom list, could be used to rapidly decipher which gloms were interconnected by a single nerve structure. This algorithm produced the same results as simply comparing all indices of the labelled edges to each corresponding index of the glom array to establish overlap, but with a processing time taking under a minute even in large arrays, as compared to hours with the rote method.

#### Creating a Master Pairwise List for Excel Export

Each sublist within the master list had its pairwise connections extracted into a new list containing every connected glom pair within the network. For example, an edge yielding the connections [10, 11, 12] to the master list would result in the pairs [10, 11] and [11, 12] being added to the ultimate pairwise list. This pairwise list was exported as a two column excel file.

#### Creating an Overlay for Labelled Glomeruli and Their Connections

To allow easy identification between glom nodes described by the 2D network and the original 3D image, the numerical identity of each labelled glom was drawn into a new tiff file. This was accomplished by identifying the centroid coordinate of each glom in the downsampled array, then drawing the labelled numerical identifier of that glom at the centroid location into a new 3D array of the same dimensions using the Python Imaging Library module. Similarly, the centroids of all glomeruli that contained pairs in the network excel export had white lines drawn between them within a seperate tiff, for connectivity visualization and validation. The resulting arrays were saved as a tiff and imported in Imaris overtop the original isosurface to visualize the relationships between the network and the 3D image. For ease of visualization, an smoothed-isosurface was applied overtop the network-connections overlay when showing large 5x networks in any figures.

#### NetTracer3D V2 Improvements

Although it was not used for the majority of the results of this study, a much higher performance search algorithm was developed in tandem with the progression of the study to be used in place of the list-array search used in V1 20x network generation for future analysis; one that allowed similar performance without any downsampling step, effectively providing a 125x increase in computational function. This method instead iterated through all labelled gloms, calculated their dilation space, before punching out a small subarray for each glom from the larger glom array. After allowing the glom to dilate in its subarray, an equivalent subarray was punched from the nerve array. The labelled edge-IDs interacted with by the glom in this space were designated as connections for that glom and stored in a dictionary with the glom label as their key. After processing all gloms in parallel for increased speed, all glom dictionaries were sorted through and shared connections were extracted as pairs to be saved in the network excel file. To compensate for the lack of any downsampled steps, this alternate algorithm allowed the edges to instead be dilated by some factor, resulting in similar logic and outcomes as the V1 algorithm, but with enhanced user-parameter control. Networks generated by V1 and V2 for the neuroglomerular networks were overall very similar, but not necessarily identical due to V1 dilating nodes in a lower-fidelity downsampled array compared to V2, which permitted more pixel-specific dilation in the full-sized array, V1 utilizing downsampling to reconnect nerve masks compared to V2 using dilation in a full-sized array, again providing V2 greater specificity, and finally V1 utilizing dilation followed by watershedding to create 5x networks, leading to the loss of a small number of node identities, compared to V2’s smart-dilation (see below) function which allowed dilation while retaining labels without losing any nodes.

#### V2 Smart-dilation

While the subarray method of V2 could also be used to generate true, overlapping search-space networks at 5x, although it lagged in performance if the number of nodes numbered in the thousands. As the array trim 5x method already used in V1 proved to be the fastest option of any of these methods, the use of this algorithm is preserved as an option in V2. Rather than the previous V1 5x method of having to rely on outside dilation followed by a second round of watershedding to reestablish Glom-IDs, V2 instead provides the user the option of using a smart-dilation algorithm. Smart-dilation allows dilation of a pre-labelled array, such as those created by MorphoLibJ, Cellpose, or ndimage, without the loss of labelled IDs. Although dilation is typically designed for binary arrays, the smart-dilate option functions by first obtaining the dilated binary output as normal, but replacing the cores of the dilated array with the original labelled array, resulting in labelled nodes with a binary shell. The indices of the binary shell were then assigned the label of its nearest labelled neighbor through the use of a distance transform, resulting in a label-preserved dilation. Network detection in NetTracer3D V2 using smart-dilation allows for the fast array trim 5x network to be performed on any labelled array, preserving performance on multi-gigabyte arrays containing nodes numbering in the tens of thousands, but did not provide the functionality of overlapping search regions.

#### Creating and Quantifying the Network

The python module ^50^was used to create a network from the pairwise excel file containing all glom-glom connections. Community detection within the network was accomplished via the networkX Louvain algorithm ^51^. Louvain was chosen as it is an efficient community detection algorithm and is well suited for “defining communities to use as clustering variables in analysis” ^52^. When generating Louvain networks, any repeated edges connecting the same pair of gloms gave that connection an additional weight in the network, with a weight of one meaning that only one neural connection between gloms was found, a weight of two representing two connections, etc. All community related quantifications utilized this weighted Louvain detection, but in instances of large, dense 5x networks, an unweighted label propagation community detection algorithm ^53^ was used solely for visualization due to its high speed.

#### 20x Network Analysis

Statistical analysis of networks was largely using the python NetworkX package. Louvain networks were quantitatively described by the networkX modularity function ^24^, obtained from Louvain community detection. Other statistics obtained directly from NetworkX methods included counts of gloms in a network, the number of connections per glom in the network, and the total number of communities detected by the Louvain algorithm. For quantitative analysis at 20x, only the largest connected component within an FOV was used to obtain a modularity score, as we were primarily interested in micro-network structure between gloms at high resolutions, meaning smaller communities at the edges of networks were excluded from this portion of analysis. High density FOVs (SK1 and SK2) were prioritized for network analysis since they provided by far the most information in a single image, but a small number of low-density FOVs were also included to demonstrate the gradient in community structure between networks containing a low and high number of gloms.

To establish significance in the community structures, all analyzed network FOVs had a random network generated of equivalent size to the network’s largest component, and containing the same number of nodes and edges. Nodes in the random network were iterated through serially and randomly assigned a partner until their network contained an equivalent number of edges as the volume they were derived from.

The random network for each volume was generated 100 times, and its modularity was averaged. This averaged, random distribution of modularities was finally compared to the distribution obtained from samples using a two-sample t-test. This method was also used to create random networks for 5x comparison. **(Supplemental Table S9)**

#### 5x Network Analysis

Creating a network out of a 5x sample with the nerve trunk present often revealed highly aggregated networks with low modularity and a non-obvious presence of communities. This was due to the majority of gloms being along a primary nerve tract that stretched through the entire volume, while the reduced resolution of 5x made assessing finer glom-nerve-glom connections unfeasible. This tract included the large nerve trunk that followed the arcuate vessel, and its upward projections alongside interlobular vessels into the cortex to seek out glomerulus vascular poles.

Since we were interested in analyzing the nature of secondary glom-nerve-glom connections, it was pertinent to also develop an analysis method that disregarded this primary trunk to reveal communities in 5x. This was accomplished by removing the trunk after breaking the nerve network into discrete glom-nerve-glom units. After the trunk was removed, all connections that remained were either those that traveled directly from glom-to-glom, resulting in a massive increase in overall network modularity. Primary trunk removal was indicated by the modularity spiking by several factors after this algorithmic step, after which analysis on the network could proceed. In two instances, the algorithmic step to remove the largest nerve structure after discretization had to be repeated as the arcuate-associated nerve was outside the FOV, meaning there were multiple “trunks” arising from large interlobular vessels. In one instance (SK18), the main nerve trunk was not removed due to being outside the volume, causing the network to have an initially high modularity. It must be noted that due to the heterogeneity of the samples, the degree to which the trunk was removed likely varied, and therefore these networks are best treated as predictions at secondary structure rather than true descriptions.

5x networks were also analyzed with Louvain detection and other associated functions from networkX. To establish the successful removal of a trunk, modularity first was calculated as an aggregate for all disconnected components, rather than only on the largest connected component. This meant treating each component as its own community in the overall network, rather than analyzing each component separately. A high spike in modularity meant secondary groups had properly split from the main trunk. Further analysis was then performed on large connected components of the resulting 5x networks. The proportion of glomerular interconnectivity was likewise obtained by dividing the number of glomeruli in the trunk network by the number in the largest component of the trunkless network, acting as an approximate descriptor of what size modules emerged with secondary connectivity after trunk removal. **(Supplemental Table S13).**

#### Performing Degree Distribution Analysis of Networks

Degree distribution analysis of both 5x and 20x networks was accomplished in Python using NetworkX. First, the node-pair excel files were used to populate a network with repeated connections assigned an additional weight. The degree of each node, representing the number of connections that node possessed, was obtained for each node from networkX and saved in excel. The excel FREQUENCY() function was used to obtain a count of the number of nodes possessing some degree k, which was converted into a proportion of nodes possessing some degree k from 1 to the maximum degree in the network. These values were used to generate a scatterplot with degree on the x axis and the proportion of nodes with that degree on y. As any degree corresponding to 0 nodes made power-curve modelling impossible, all proportions were incremented by 0.001 prior to modelling. **(Supplemental Table S10).**

#### Isolating Large 5x Networks After Removing Nerve Trunk

After removing the nerve trunk in applicable volumes, 5x networks often revealed a nerve network connecting the majority of gloms with strong community structure. This interconnected network and its associated nerves were isolated with Python. NetworkX was used to generate a list of all glom IDs in the target connected network component, as identified by the networks script. The numpy function isin() was then used to create a binary mask from all interconnected gloms contained within this network, which was saved as a tiff and exported on top of the Imaris isosurface. Nerve connections between these gloms were isolated by converting the interconnected glom network mask to a boolean array and multiplying it against the labelled, dilated edge array. Edge IDs that remained were compiled in a list. The numpy function isin() could then be used to remove all nerves with IDs not present in this list, and the isolated nerve array was saved as a tiff to be imported into Imaris.

#### 5x vs 20x Networks

By isolating subnetworks of gloms, we used the 5x-20x coregistration to compare network prediction ability of gloms between 5x and 20x volumes.

Due to the lower resolution of 5x volumes, networks generated were expectedly less complex than the true nerve structure. For example, one hourglass structure rated by the networks script and easily visible at 20x in sample SK2 FOV 13 was seen to be broken into two disconnected communities in the corresponding SK2 5x network. As demonstrated by this observation, 5x networks lost connectivity both within and between communities, and therefore were best suited for finding high-throughput mother gloms and appraising a simplified, macro community structure.

#### Segmenting and Analyzing Confocal Immunofluorescence Microscopy Datasets

Analysis of confocal microscopy datasets was done with the same strategy as the 3D automated 20x innervation, where a skeletonized nerve segmentation was quantified within a designated region surrounding a segmented glomerulus mask. A labkit model for nerves was trained on the confocal datasets. To eliminate signal-intensity based bias, the diabetic and reference confocal datasets were first stacked together and used to simultaneously train one model. This model was used to segment all nerves, with the resulting mask being skeletonized and appraised for skeleton density around glomeruli in Python. **(Supplemental Table S14).**

#### Preforming Statistical Testing and Visualization

All dataset visualization was done with Imaris, while graphical visualization was done with matplotlib or Microsoft Excel, and figures and movies were assembled in adobe illustrator and adobe premiere pro, respectively. All statistical testing was done using the scipy stats module. Data were tested as shown with the exception of radial angles, for which all angles were transposed away from radial arrangement by adding 6.28 to any angles below 2 radians during script execution. This was because of the circular nature of radial distributions, where an item of 2 radians in actuality is only 2 radians removed from one of 6.28, rather than 4.

For the purposes of this data, moving all low-value radian datapoints to the top of a now-linear distribution allowed the entire dataset to be statistically tested without being incorrectly appraised due to the radial nature of the distribution.

## Software versions

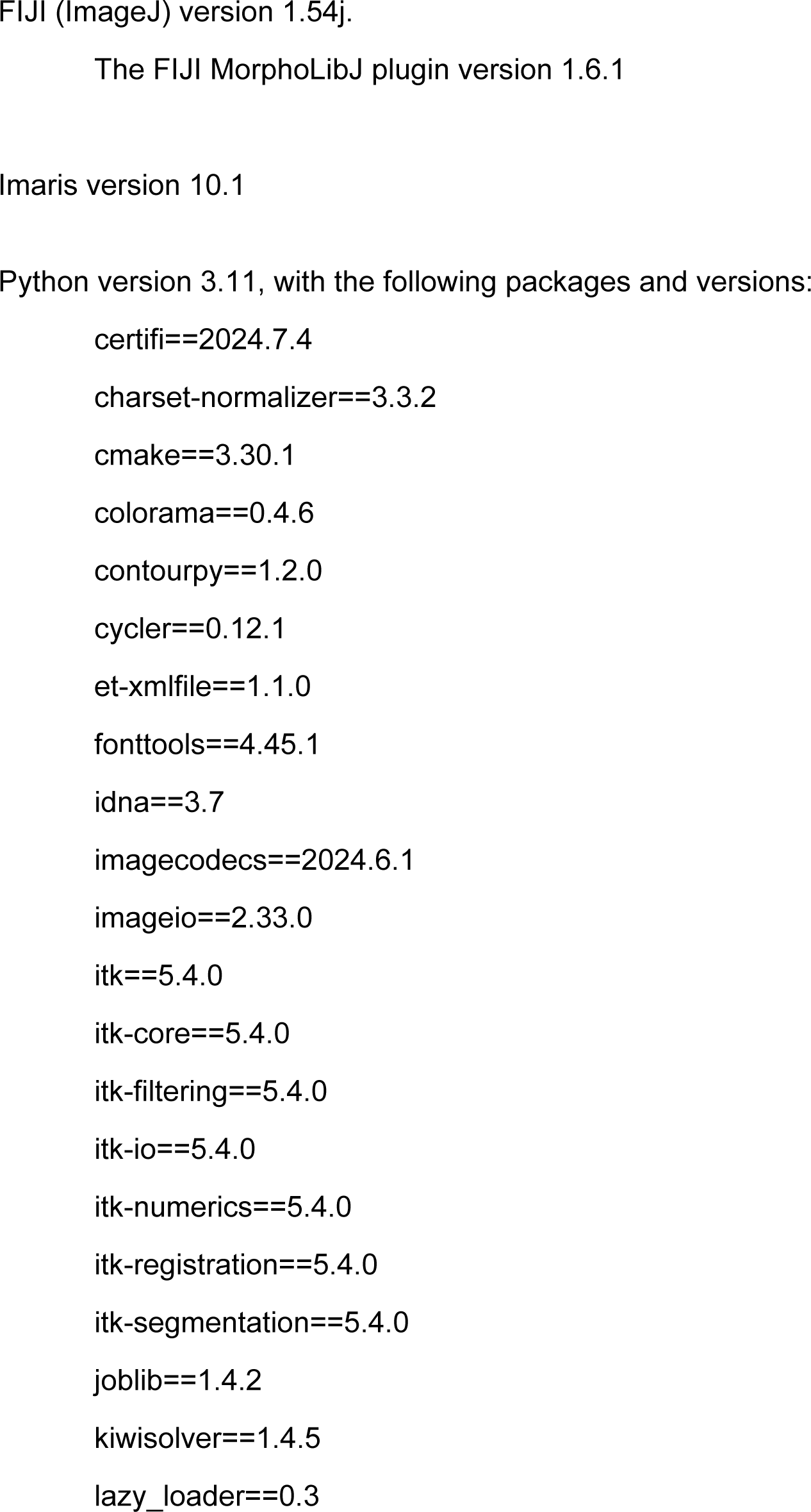
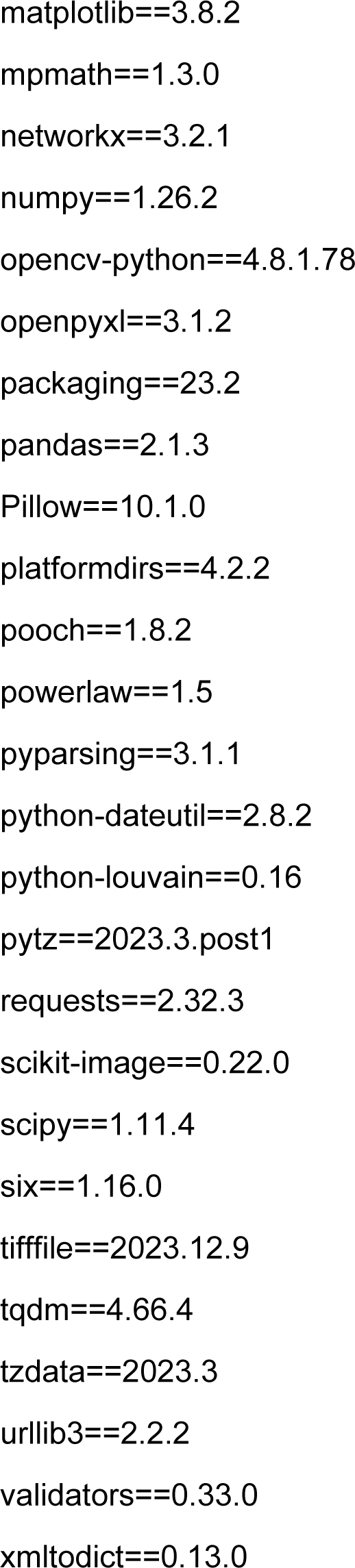

## Supporting information

Supplemental tables for the main manuscript

## Acknowledgements

We thank Kristy Conlon for recruiting patients and assistance with sample processing. Pediatric donor kidney tissue was supplied through the United Network of Organ Sharing, Organ Procurement and Transplantation network and the International Institute for Advancement of Medicine, organizations that link donor families to the scientific community. We are very grateful to the families who generously donated such precious gifts to support research. Imaging/Analysis with the Zeiss Lightsheet 7 and Imaris were performed in part through the use of Washington University Center for Cellular Imaging (WUCCI) supported by Washington University School of Medicine, The Children’s Discovery Institute of Washington University and St. Louis Children’s Hospital (CDI-CORE-2015-505 and CDI-CORE-2019-813) and the Foundation for Barnes-Jewish Hospital (3770 and 4642). We are grateful to the Human BioMolecular Atlas Program (HuBMAP) consortia for supporting this project and providing resources for data sharing. We thank Praveen Krishnamoorthy and Peter Bayguinov of the Washington University Center for Cellular Imaging (WUCCI) for their advice, support and resources in helping implement the light sheet clearing and imaging protocol and Masato Hoshi for initial experiments in evaluating LSFM method. We thank the Washington University Kidney Translational Research Center (KTRC) supported in part by the Division of Nephrology for the infrastructure for human tissue processing, storage and specimen tracking. We thank our colleagues from the HuBMAP-TMC, Tarek M. El-Achkar, Seth Winfree, Michael T. Eadon and Lingyan Shi for their guidance and HuBMAP-HIVE teams especially Nils Gehlenborg and Philip Blood for their efforts in visualization and data accessibility of LSFM images. This work was in part supported by National Heart, Lung, and Blood Institute (NHLBI) Molecular Atlas of Lung Development Program, Human Tissue Core (LungMAP HTC) (U01HL148861 to G.S. P.), HuBMAP (U54DK134301 to S.J.) and Pediatric Center of Excellence in Nephrology at Washington University (P50DK133943 to S.J.). The content is solely the responsibility of the authors and does not necessarily represent the official views of the National Institutes of Health.

## Author Contributions

**S.J.** conceived the idea and procured funding. **L.M.** wrote the first draft and designed the figures/movies. **S.J., L.M.** finished the final draft. **A.L.K.** collected metadata, prepared adult tissue, performed histology and experiments. **J.P., H.H**. and **G.S.P**. developed methods to procure kidney tissue from donor sites and prepared pediatric kidney tissue for LSFM experiments. **B.Z.** developed the LSFM immunostaining procedure and performed all the LSFM and confocal microscopy validation experiments. **M.K.** provided project management and data organization and deposition. **L.M.** designed pipelines for and carried out the LSFM imaging and analysis, and wrote the code. **S.S.** performed mathematical modeling and helped conceive analytical pipelines. **J.P.G**. and **S.J**. evaluated gross and microscopic pathology. All authors reviewed and edited the manuscripts.

## Data Availability

All 3D LSFM image files and processed segmentation masks will be available from HuBMAP consortium (www.hubmap.org) after publication. The source code for analysis will be available from a public Github repository accessible through a peer-reviewed publication. All movies are available through Zenodo https://doi.org/10.5281/zenodo.12802740 (DOI:**10.5281/zenodo.12802740)**.

## Code Availability

The code used for analysis will be available through a Github repository upon publication. Note that NetTracer3D tools for network analysis is freely available for academic and nonprofit use, provided that this manuscript is cited in manuscripts or presentations utilizing NetTracer3D. Commercial use is available for a fee. Copyright © is held by Washington University. Please direct all commercial requests for licensing, information, and limited evaluation copies to Washington University’s Office of Technology Management at.

## Competing Interests

S.J. and L.M. have an intellectual property invention disclosure on the 3D nerve network analysis in solid organs software tool NetTracer3D the copyright of which is held by Washington University in St. Louis and may receive royalties from commercial use.

## Information for Data Requests and Correspondence

Sanjay Jain

## Supplementary Items

### Supplementary tables

Supplemental Table S1. Sample information and markers used for light sheet fluorescence microscopy.

Supplemental Table S2. Sample staining information and quality determining use in quality and quantitative assessment. Structures and cell types.

Supplemental Table S3. Data for polar and azimuth angles of glomeruli measured at various depths from 20x FOVs from 5 samples at varying cortical depth.

Supplemental Table S4. Longitudinal Distribution of Glomeruli in Cortex. Supplemental Table S5. Longitudinal Distribution of Collecting Duct.

Supplemental Table S6. Longitudinal Distribution of Glomerular Innervation Density.

Supplemental Table S7. Automated Nerve Innervation Scores for various 20x FOVs across Several Samples.

Supplemental Table S8. Manual Nerve Innervation Densities for various 20x FOVs across Several Samples.

Supplemental Table S9. Modularities of 20x Networks.

Supplementary Table S10. Contains the proportion of nodes with degree k for several 5x networks, used to construct a degree distribution by modelling a power curve.

Supplemental Table S11. Includes a variety of statistics about 5x images, including sample volumes, glomerulus count, cortical glomerulus density, proportion of sample that is cortex, and the average innervation score of glomeruli in the 5x image.

Supplemental Table S12. Morphological stats about glomeruli at 20x resolution.

Supplemental Table S13. Information about 5x Networks.

Supplemental Table S14. Glomerular innervation scores of 2D confocal immunofluorescence microscopy sections.

Supplemental Table S15. Glomerular Cell Counts and Densities.

Supplemental Table S16. Information about FOV coregistrations for various samples.

### Extended Supplemental Data Figures

Supplemental Figure S1. Supporting Images for Figure 2: 3D LSFM.

Supplemental Figure S2. Supporting Images for Figure 4: Characterization of neuroglomerular networks at both 5x macro and 20x micro scales.

Supplemental Figure S3. Supporting Images for Figure 6: Altered neuroglomerular networks and morphology with age and disease using LSFM.

### Movies (available to view in Zenodo DOI: 10.5281/zenodo.12802740)

**Supplemental Movie 1:** 3D view of the entire slice showing key structures.

**Supplemental Movie 2:** Relationship of nerves with glomeruli and juxtaglomerular apparatus.

**Supplemental Movie 3:** Neuro-nephron connectivity.

**Supplemental Movie 4:** Neurovascular – nephron patterns in the medulla.

**Supplemental Movie 5:** Network motifs.

**Supplemental Movie 6:** LSFM movie of pediatric kidney

**Supplemental Movie 7:** Neuronephron connectivity time course

## Movie Legends

Movie 1: 3D view of the entire slice showing key structures.

3D light sheet fluorescence microscopy 5x movie of reference adult sample SK3, demonstrating glomeruli, collecting ducts, nerves, and blood vessels. 0:00s — Raw signal. 0:10s — Segmentations. Annotations are in the movie.

Movie 2: Relationship of nerves with glomeruli and juxtaglomerular apparatus.

The movie depicts innervation of glomeruli in 2D optical sections, containing glomeruli, Tuj1(labels TUBB3)- stained nerves, and CGRP-stained sensory nerves. 0:22s — Innervation of the JGA. 0:33 s— Innervation of the Macula Densa. 0:47s — Innervation of the outer boundary of the Bowman’s Capsules.

Movie 3: Neuro-nephron connectivity.

The movie explores innervation between different structures of the same nephron, and between nephrons in both 3D and 2D optical sections, containing glomeruli, Tuj1 (TUBB3)-stained nerves, CGRP-stained sensory nerves, proximal (convoluted) tubule, thick ascending limb, distal convoluted tubule, and collecting duct. 0:00-1:53min — 3D relationships. 0:38s — Innervation of glomerulus JGA. 1:02min — Post-JGA innervation of medullary ray structures. 1:54min-end — 2D relationships. 2:38min — Interglomerular/internephron innervation.

Movie 4: Neurovascular – nephron patterns in the medulla.

The movie shows innervation pattern within the medulla. 0:00s—5x adult medullary innervation pattern in 3D; 0:34s—in 2D also showing Vasa Recta and Collecting Duct; 0:48s—in 3D at 20x resolution. 1:04min—20x adult medullary innervation of proximal tubule, thick ascending limb, and Vasa Recta in 3D; 1:31min—view if the previous in 2D. 2:11min—5x adult medullary innervation pattern in 3D; 2:40min—in 2D also showing Vasa Recta and Collecting Duct; 2:54min—in 3D at 20x resolution; 3:15—in 2D at 20x resolution.

Movie 5: Network motifs.

Exploring 3D neuroglomerular networks at 5x and 20x resolution. 0:00s — Raw 5x signal from young adult sample SK2. 0:16s— Segmented 5x SK2 with 20x coregistrations. 0:26 — Exploring SK2 20x FOV. 0:38 — 20x network featuring hourglass motif. 1:19 5x “Type I” network in SK2. 1:41 — Segmented 5x adult SK3 sample featuring a “Type 2” network containing a lattice motif. 2:23 — Exploring SK3 20x FOV, featuring a network with a lattice motif.

Movie 6: LSFM movie of pediatric kidney

3D lightsheet 5x image of neonatal sample SK414, containing glomeruli, collecting ducts, nerves, and blood vessels. 0:00s — Raw signal. 0:22s — Segmentations. 1:34min — Overlayed segmentations.

Movie 7: Neuronephron connectivity time course

Exploring neuronephro-networks across a life time course in 1mm^3^ 20x images. 0:00 — Raw neonatal. 0:17sec — Segmented neonatal. 0:47 sec— Raw infant. 0:54 — Segmented infant with network. 1:13min — Raw young adult. 1:23min — Segmented young adult with network. 1:33min — Raw adult. 1:43min — Segmented adult featuring network with keychain motif. 1:49min — Raw aged. 1:59min — Segmented aged with network featuring pyramid motif.

---

**Supplemental Figure S1.**
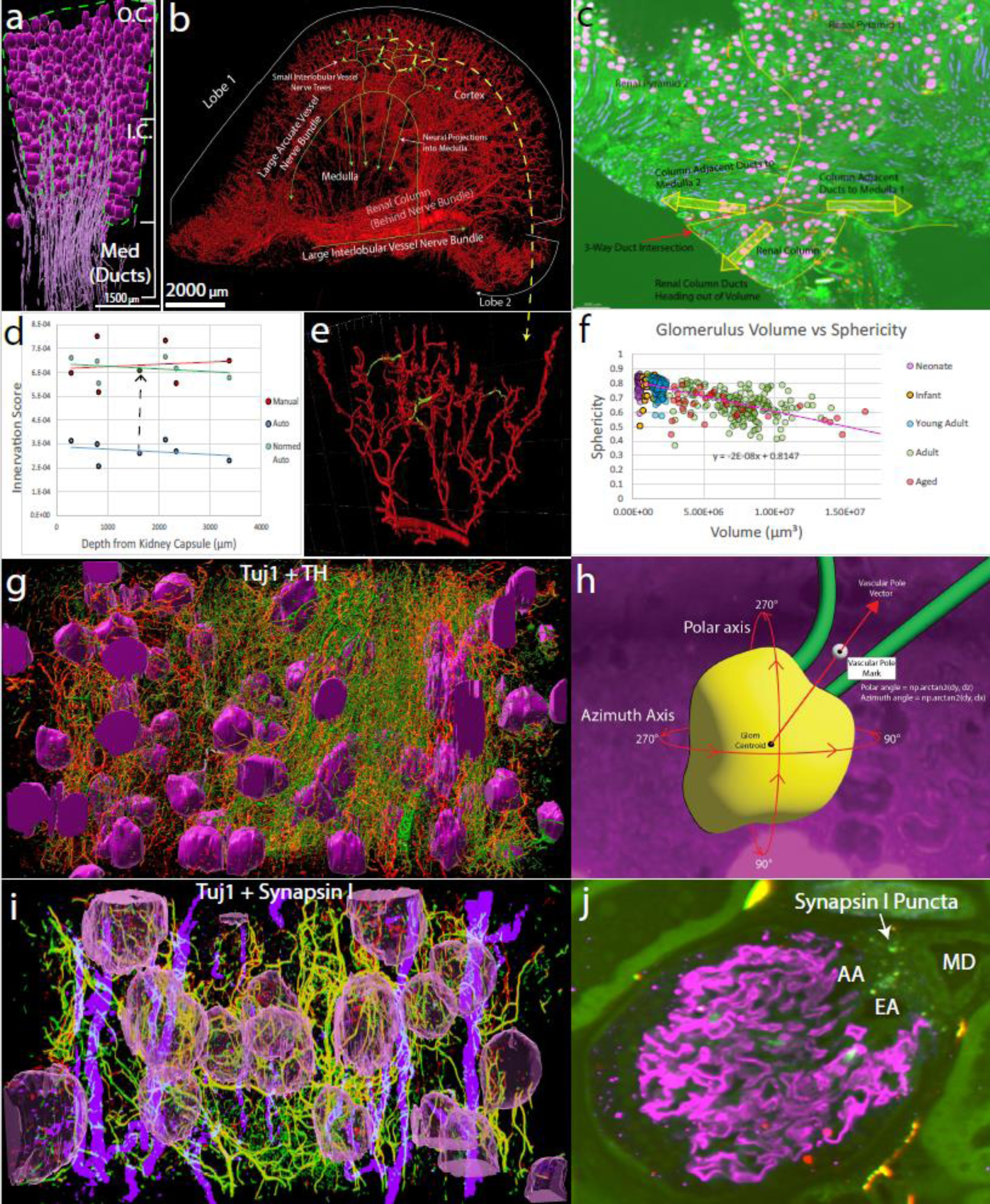
**Supporting Images for Figure 2: 3D LSFM. A.** Young adult segmented 5x image (sample SK2) with glomeruli (magenta) and collecting duct (purple). An overlay has been applied over the image to illustrate the relationship between glomeruli columns in inner cortex (IC) and glomeruli tufts in the outer cortex (OC), with medullary rays converging towards the medulla (Med). **B.** Segmented nerves from an entire 5x renal lobe of a pediatric patient (sample SK14) demonstrating the dense neural arborization of the cortex and other kidney regions. Nerve trees in the cortex emanate in as separate branches from the arcuate-artery associated nerve bundle, but reconnect a certain distance into the cortex. **C.** 2D view of the renal column from 5x pediatric patient SK14 showing the divergence point (‘3-Way Duct Intersection’ – Circled red) for collecting ducts (purple) heading into either adjacent medulla, or down the renal column. Yellow arrows show the CD directions. The renal column itself is innervated by smaller vessels that emanate from the large interlobular vessel tracking up towards the arcuate arteries. **D.** Validation of automated neural analysis method by comparing neural innervation to glomerular depth from kidney capsule (µm). Nerve lengths within 10 µm of 120 glomeruli were manually counted across seven FOVs in samples SK1 and SK2 (PPID: 3785), with the nerve length µm/µm^3^ graphed in red. The same FOVs were automatically appraised, representing ∼194 glomeruli, are graphed in blue, with the points representing the density of segmented nerve-skeleton pixels within 10 µm of glomeruli. These distributions both show shallow trends indicating innervation of glomeruli is similar between cortical layers and supports that the automated analysis is a reasonable surrogate measurement of innervation magnitude around an object, albeit not directly comparable to manually-measured nerve length. The automated analysis was normalized to be similar in overall magnitude to the manual measurements (by adding to all points the difference indicated by the dotted black line), graphed in orange with a black outline, with the normalized automated and manual distributions not being significantly different by Welch’s t-test (p=0.761). Note that the manual measurements in an FOV only included glomeruli completely included in the volume while the automated measurements used all glomeruli, which may explain some visible differences. **E.** A closer view of the cortical neural trees from the segmentation in B, clearly demonstrating how separate neural trees reconnect in higher cortical regions (denoted by nerve segments colored green). **F.** A graph comparing the volumes and sphericities of 643 glomeruli measured in this study, demonstrating a downward trend where glomeruli lose their spheroid shape as they enlarge. Data points are color coded by age group. Correspondingly, glomeruli appear most spherical in younger adult patients with smaller glomeruli. **G.** A segmented image (magenta - glomeruli, red filaments - Tuj1 (TUBB3) stained nerves, green filaments - TH-stained sympathetic nerves) from sample SK24 demonstrating how sympathetic nerve stain appears to ubiquitously overlap with general nerve stain. **H.** A cartoon depicting how glomeruli angles were calculated. The centroid of the glomerulus was used to generate a vector to a point placed to mark the vascular pole, which could be used to calculate angles with the numpy arctan2() function. **I.** A segmented image (magenta glomeruli, red filaments - Tuj1 stained nerves, green filaments – Synapsin I-stained nerves) from sample SK18 demonstrating how Synapsin I stain appears to ubiquitously overlap with general nerve stain, suggesting synapses line all neural projections within the kidney. **J.** A 2D slice from the volume shown in I depicting a glomerulus and its juxtaglomerular apparatus (JGA). Synapsin I punctata (green) are abundant at the JGA, between the MD and arterioles, indicating a high density of synaptic targets in this region.

**Supplemental Figure S2.**
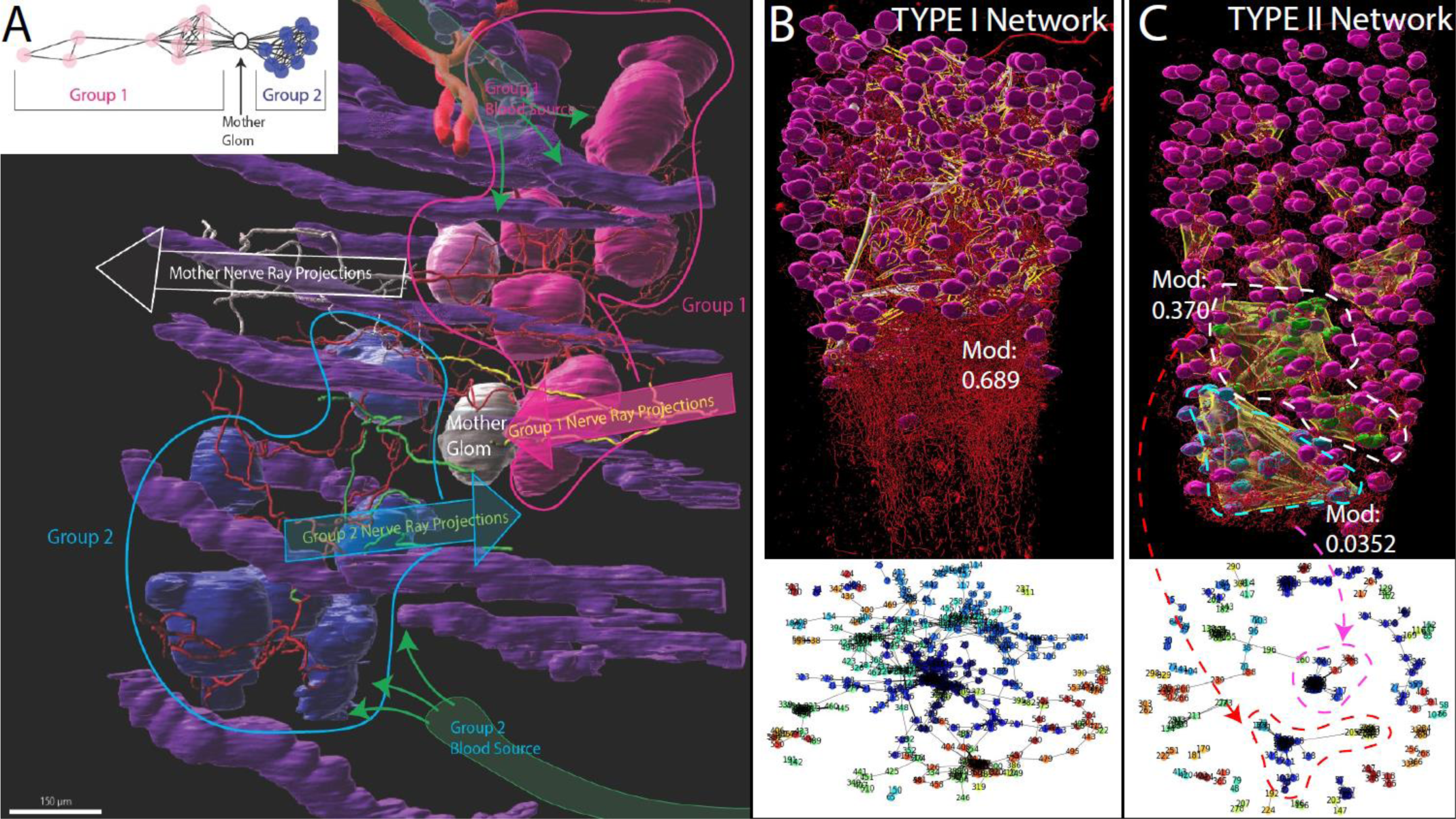
Supporting Images for Figure 4: **Characterization of neuroglomerular networks at both 5x macro and 20x micro scales. A.** Detailed view of the hourglass motif in the main text Figure 3F. Spheroid glomeruli are surrounded by manually segmented nerves. Separate communities (blue and magenta) are unified by white mother glomerulus (glom), reflecting the network in the top left. The blue glomeruli are vascularized by an interlobular vessel from below, as annotated, while the magenta/mother glomeruli are vascularized from a distinct interlobular vessel from above. Neural projections from the blue community are colored green and denoted by the blue arrow, while magenta community ray projections are colored yellow and denoted by the magenta arrow. Ray projections appearing to come from the ‘mother glomeruli area’ are colored white, denoted by the white arrow. Inset, schematic representation of the network. **B.** Segmented glomeruli (magenta) and nerves (red) showing an example of a type I 5x network, obtained from sample SK1 (PPID: 3785), with a network overlay in yellow/white and its network shown below. Type 1 networks are defined by their high connectivity, containing most glomeruli in a single connected, modular network component. 81% of all glomeruli in the non-trunk removed network are retained in a large connected component (n=365) which itself has a high modularity of 0.689. **C.** Segmented glomeruli (magenta) and nerves (red) showing an example of a type II 5x network, obtained from sample SK18 (PPID: 3785), with a network overlay in yellow/white and its network shown below. Type II networks are defined by lower connectivity, with a greater number of small modules that are unconnected beyond the main trunk. Remaining large modules themselves may demonstrate module community structure, such as the hourglass-motif circled in white (n=40, mod=0.370) while other large modules were clumpier, such as the grape-motif circled in cyan (n=36, mod=0.0352). Note that sample SK18 did not require trunk-removal option in network analysis due to the nerve trunk existing largely outside the dissected sample imaged.

**Supplemental Figure S3.**
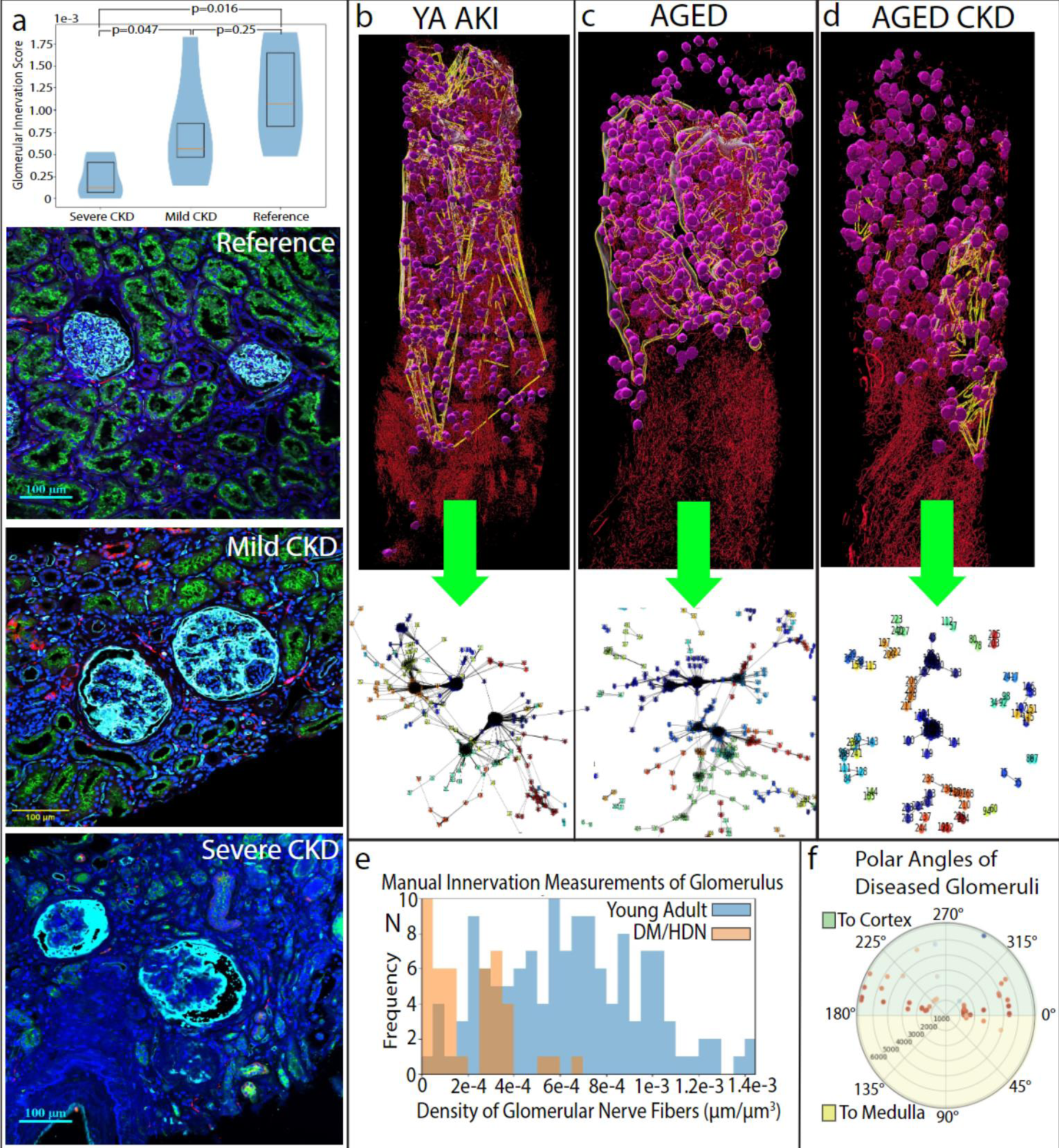
Supporting Images for Figure 6: **Altered neuroglomerular networks and morphology with age and disease using LSFM. A.** A bar graph comparing automated innervation measurements around glomeruli from sections used in 2D confocal immunofluorescence microscopy from severe CKD diabetic patients (n = 5, severe CKD Stage 4 or greater; Average 2.27e-3; SE 1.02e-4), mild CKD diabetic patients (n = 5, CKD Stage 2; Average 7.73e-3; SE 2.86e-4), and reference (n = 5, Average 1.29e-2; SE 2.59e-4) patients. Both mild CKD (p=0.047) and reference (p=0.016) were significantly greater than severe CKD with a rank-sums test, but mild CKD diabetics and reference were not significantly different from each other (p=0.25). Representative confocal microscopy images of these 3 patient groups are shown below the graph with glomeruli as white spherical structures, nerves in red, proximal tubules in green, and nuclei in blue. **B.** Segmentation LSFM images of a young adult patient (YA) with AKI (SK28; PPID: 4125) with glomeruli in magenta and sympathetic nerves in red. A network overlay is shown in yellow/white, yielding a type-1 network appropriate for their age group (young adult) and glomerular density. The network is shown below, depicting a high modularity (0.704) central structure containing the vast majority of glomeruli. **C.** Segmentation of a >75 yo aged reference patient (SK12; PPID 3900) with glomeruli in magenta, nerves in red, and network overlay in white/yellow. Below is its type-1 network containing a large central structure with the majority of glomeruli and high modularity (0.683). This sample was the only type-1 network obtainable from a patient older than the young adult group. **D.** Segmentation of an >75yo aged reference patient (SK23; PPID: 3990) with glomeruli in magenta, sympathetic nerves in red, and network overlay in white/yellow. Below is its type-2 network, with the largest structure being less modular (0.0473) than similarly sized structures in other type-2 networks. **E.** Manually measured glomerular innervation density histograms for all nerve fibers within 10 µm of glomeruli in both reference (SK1, 2; PPID: 3785) and diabetic/hydronephrosis diseased patients (SK8; PPID: 3909. SK9/10; PPID: 3702). A Welch’s t-test demonstrated reference glomeruli (n=133) receive highly significantly greater nerves than the diseased group (n=42; p=4.48e-26). **F.** Glomerular polar angles from samples SK8 and SK9 demonstrating the same two-quadrant, bimodal bias in diseased samples as compared to young adult reference samples. The radial axis is cortical depth in µm.

## Notes

https://doi.org/10.5281/zenodo.12802740

